# Whole-genome sequencing of Chinese native goat offers biological insights into cashmere fiber formation

**DOI:** 10.1101/2021.11.06.467539

**Authors:** Hu Han, Man-Man Yang, Jiang Dan, Xing-Ju Zhang, Qiang Wei, Tao Chen, Qi-Ju Wang, Cheng-Ye Yang, Bater Wulan, Ting-Ting Zhang, Gang Gen, Mengkedala, Bin Li, Wei-Dong Deng, Ze-Pu Miao, Ran Wang, Qing-Feng Zhang, Lin Li, Sheng-Yu Chao, Ming Fang, Yong Li

## Abstract

Cashmere evolved naturally in the goat, and almost all breeds of goat can produce more or less cashmere fibers. However, the genetic alterations underlying cashmere trait selection are still unclear.We sequenced 120 Chinese native goat including two cashmere goat breeds (Ujumain, Chaidamu) and six ordinary goat breeds (Jining Gray, Matou, Guizhou Black, Jintang Black, Yunnan Black Bone, Chengdu Brown). The genome-wide selective sweep of cashmere goat and ordinary goat revealed a novel set of candidate genes as well as pathways, such as Nuclear factor kappa-B and Wnt Signaling pathways. Of them, the *LHX2* gene regulating hair follicle development, was evident from the strongest selection signal when comparing the Uhumqin cashmere goat and ordinary goat. Interestingly, we identified a 582bp deletion at 367 kb upstream of *LHX2* with higher frequency in cashmere goats and their ancient relatives. This mutation probably rises along the breeding procedures, and is putatively responsible for cashmere production and diameter, as revealed by association studies. Luciferase assay shows that the deletion, which acts as an insulator, restrains the expression of *LHX2* by interfering its upstream enhancers.Our study discovers a novel insulator of the *LHX2* involved in regulating cashmere production and diameter, which would be beneficial to understanding hair follicle development and regeneration. Our findings also provide new insights into the genetic formation of cashmere, and facilitate subsequent molecular breeding for cashmere goat improvement.

## Introduction

Cashmere contributes high economic value to the textile industry. Cashmere is made from the processing of the underwool of goats that are grown in high and cold regions of the world, such as the Tibetan plateau and Mongolia. China is the largest cashmere producer globally, accounting for approximately 75% of the world’s supply(WaldronBrown and Komarek 2014). Improvement in the quantity and quality of cashmere is an important breeding goal in goat farming.

The cashmere goats with a long fine underwool, share a common ancestor with other ordinary goats that were domesticated from wild goat (bezoar) approximately 11,000 years ago in the Fertile Crescent of southwest Asia and adjacent areas(Zeder 2008; Daly et al. 2018). Almost all the breeds of goats can produce more or less cashmere fibers, but only the breeds with sufficiently fine hair are called cashmere goats (cashmere goats in China produce 250 g to 500g of fiber, while only 50g in ordinary goats)(Ryder 1993). Previous research postulated that cashmere goats would not be domesticated locally, except several highland regions including Himalayas, Mongolia, and Kirghizia(Millar 1986). This kind of domestication leads to selecting more and more cashmere wool to meet textile demands. Therefore, in contrast to cashmere fiber formation, cashmere production was pursued by intense artificial selection pressure during the domestication of cashmere goats(Li et al. 2017; Cai et al. 2020; Jin et al. 2020).

Previous transcriptome studies in cashmere goats have already found many important pathways that are involved in cashmere fiber development, including *WNT, FGF5, BMP, TGF-β, NOTCH, SHH*, and some of them may play important roles in secondary hair follicle development, such as *LHX2, FGF5* and *TGF-β4*(GengYuan and Chen 2013; Geng et al. 2014; Gao et al. 2016; Su et al. 2018; Zhang et al. 2019; Dai et al. 2020; Zhang et al. 2020). Meanwhile, some genes, such as *FGF5, SGK3, IGFBP7, OXTR, ROCK1, LHX2, FGF9, PRDM6* and *WNT2* were intensively selected in cashmere goats(Wang et al. 2016; Li et al. 2017; Zhang et al. 2018). Recent studies have identified several causal genes for cashmere growth trait, including *FGF5* and *EDA2R*(Cai et al. 2020; Guo et al. 2020). *FGF5* is a regulator of the hair growth cycle, and the disruption of *FGF5* is associated with a long hair characteristic(Wang et al. 2016; Li et al. 2019). Interestingly, the enhancer-absent *FGF5* was only found in domesticated goats rather than in the wild goats, indicating this causative mutation occurred in domestication process(Cai et al. 2020; Li et al. 2020). However, more researches are necessary to elucidate the genetic basis of cashmere fiber formation in goats after divergence from sheep.

Here, we generated genomic data from 42 cashmere goats and 78 ordinary goats in China and conducted comprehensive population genomic analyses. Integrated with goat genomes, we identified genetic footprints under artificial selection during goat migration and domestication. When compared the ancient and wild goat, we also investigate the origin and biological function of causative mutation for cashmere fiber formation.

## Results

### Population sequencing and genetic diversity

A total of 120 domestic goats representing eight geographically diverse breeds in China were selected for genome resequencing (Supplementary Table 1). These datasets were also analyzed together with the published whole-genome sequencing (WGS) datasets of 116 individuals from five breeds (Supplementary Table 2 and 3). Among the ten domestic goat breeds, Liaoning cashmere goat (LNC), Inner Mongolia cashmere goat (IMC), Ujumqin cashmere goat (UC) and Chaidamu cashmere goat (CDMC) are the cashmere goat breeds located in the north of China; Chengdu brown goat (CDB), Guizhou Black goat (GZB), Yunnan black bone goat (YNBB) and Jintang black goat (JTB) in the southwest and Matou Goat (MT) and Jining gray goat (JNG) in the middle are non-cashmere goat breeds (Fig. 1a). The remaining goat breeds include Korea goat (KO), Iranian wild goat (IRW) and Angora goat (ANG) from Korea, Iran and France, respectively.

**Fig. 1.**
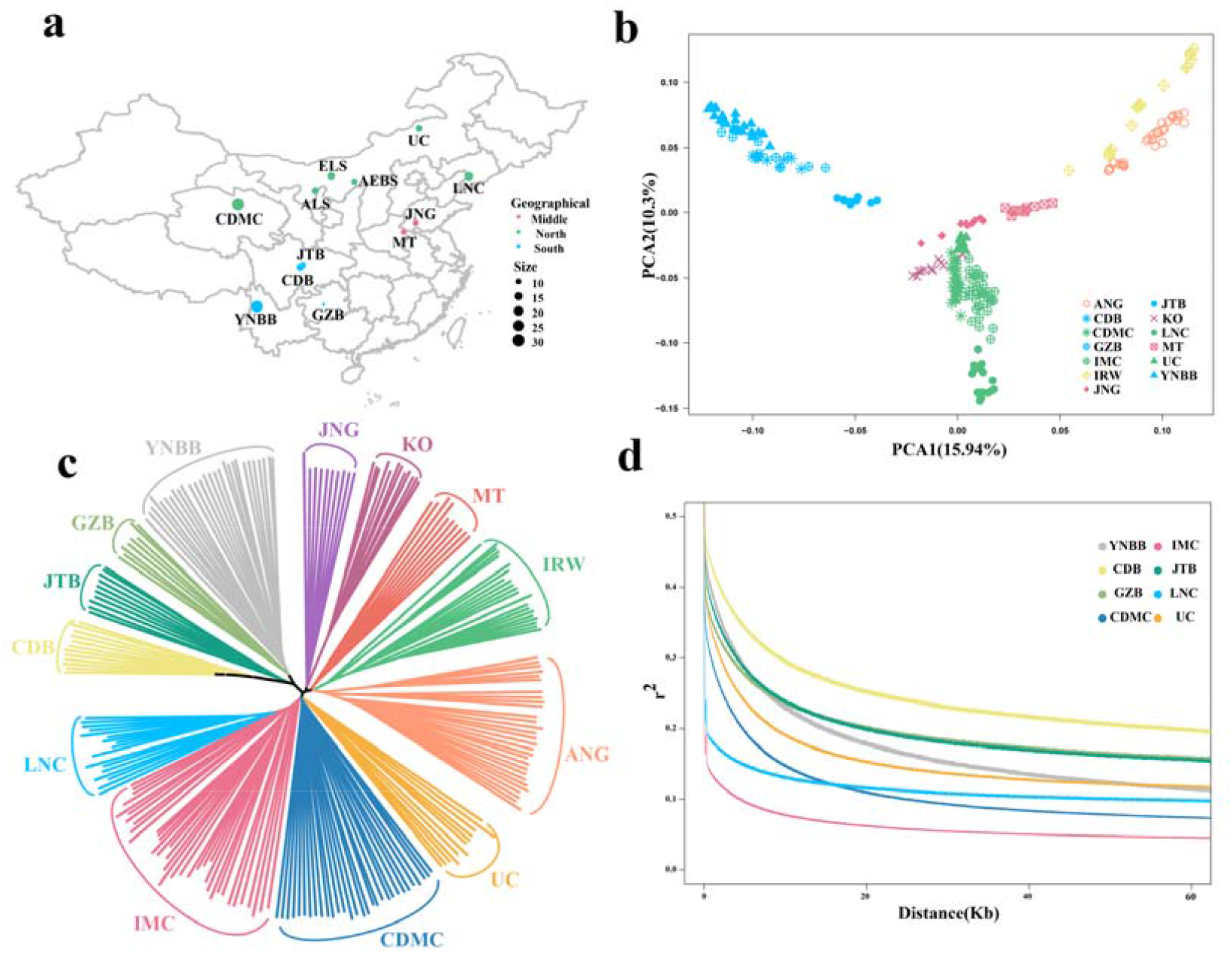
Geographic distribution and population genetic analyses of 13 goat breeds. Geographical distribution of domestic goat breeds in China. Green, blue, red represents northern, southwest, middle regions, respectively. **(b)** Principal component (PC) plot of the first two components. The fraction of the variance explained is 15.94% and 10.3% for PC1 and PC2, respectively. **(c)** Neighbor-joining tree constructed using p-distances between different breeds, including LNC (Liaoning cashmere), IMC (Inner Mongolia cashmere), UC (Ujumqin cashmere), CDMC (Chaidamu cashmere), CDB (Chengdu brown), GZB (Guizhou Black), YNBB (Yunnan black bone), JTB (Jintang black), MT (Matou), JNG (Jining gray), KO (Korean), IRW (Iranian wild), and ANG (Angora). **(d)** Linkage disequilibrium decay of goat populations measured by r^2^.

A total of 8448.92□Gb paired-end DNA sequence data were obtained from 236 goats. All goats were sequenced with an average 12-fold depth of the genome (4.17∼29.9×) and the average genome coverage of 97.82% (Supplementary Tables 1-3). 99.2% reads were mapped to the latest goat reference genome ARS1(GCA_001704415.1) (Supplementary Tables 1-3). We totally detected 13,069,924 high-quality single nucleotide polymorphisms (SNPs), with 0.605% (0.229 million) located in exonic regions (Supplementary Table s10). The average transition-to-transversion (Ti/Tv) ratio was 2.46 for all goat samples, which indicated relatively low potential random sequencing errors (Supplementary Table s10).

The principal component analysis (PCA) of the 236 goats revealed genetically distinct clusters according to their geographic locations. The clustering results of the northern cashmere goat populations (IMC, LNC, UC and CDMC) and ordinary goat populations (YNBB, JTB, GZB and CDB) were clearly separated (Fig. 1b). JNG and MT were divided into subgroups between cashmere goats and the ordinary southwest goats. Samples of Iranian wild goats (IRW) and Angora goats (ANG) (France, South Africa, and Madagascar) were significantly different from those of Chinese native goats (Fig. 1b). This result was confirmed by the phylogenetic tree using the same SNPs (Fig. 1c). In the analysis of linkage disequilibrium (LD), four ordinary goat breeds from southwest China (YNBB, JTB, GZB and CDB) showed an overall slower decay rate and a higher level of LD than the cashmere breeds from northern China (IMC, LNC, UC and CDMC) (Fig. 1d).

In the STRUCTURE analysis, When K = 4, we observed five separate clusters: IRWG and ANG in west Asia, YNBB, GZB, JTB and CDB in southwest China; cashmere goat in north China; MT and JNG in mddle east China; and Korean goats in south Korea. At K = 6, goats in the southwest China further split into two geographic subgroups: the Yunnan-Kweichow Plateau group including YNBB and GZB goats, and the Chengdu Plain group including CDM and JTB goats. Two west Asian goats (IRW and ANG) were also separated (Fig. 2a). Some cashmere goats showed evidence of admixture, which may be attributable to shared ancestral polymorphism and recent introgression events by crossbreeding with neighboring domestic goats. Interestingly, the genetic structure of the Chinese goat population revealed that UC and LNC goats represent two different types, and other cashmere goats are a mixture of these two types (Supplementary Fig. 3).

**Fig. 2.**
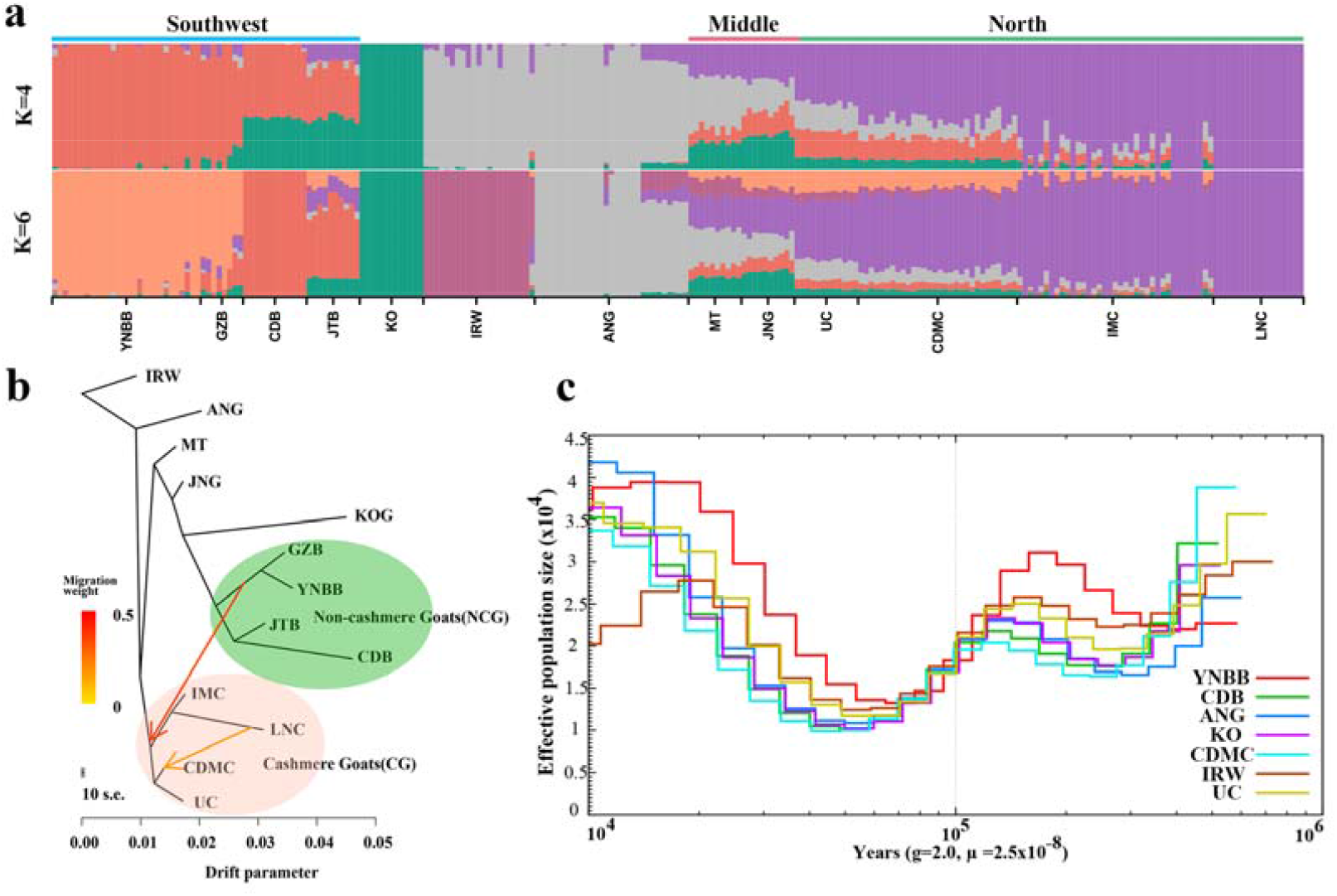
Population structure, gene flows, and effective population size of goat populations. **(a)**Model-based population assignment with ADMIXTURE analysis for K = 4 and 6, respectively. The population names are at the bottom of the figure, and the geographic locations are at the top. Phylogenetic network of the inferred relationships among the 13 native breeds with two inter-group migration edges being identified. The branch length is proportional to the drift of each population. IRW (Iranian wild goat) was used as the outgroup to root the tree. The colored regions in the phylogenetic tree represent two inferred genetic groups. Arrows indicate migration events, and a spectrum of heat colors indicates the migration weights of the migration events. **(c)** Pairwise sequentially Markovian coalescent analysis for the representative individuals sequenced at a high read coverage, exhibiting inferred variations in Ne over the last 10^6^ years. g (generation time) = 2 years; μ (neutral mutation rate per generation) = 2.5× 10-8.

**Fig. 3.**
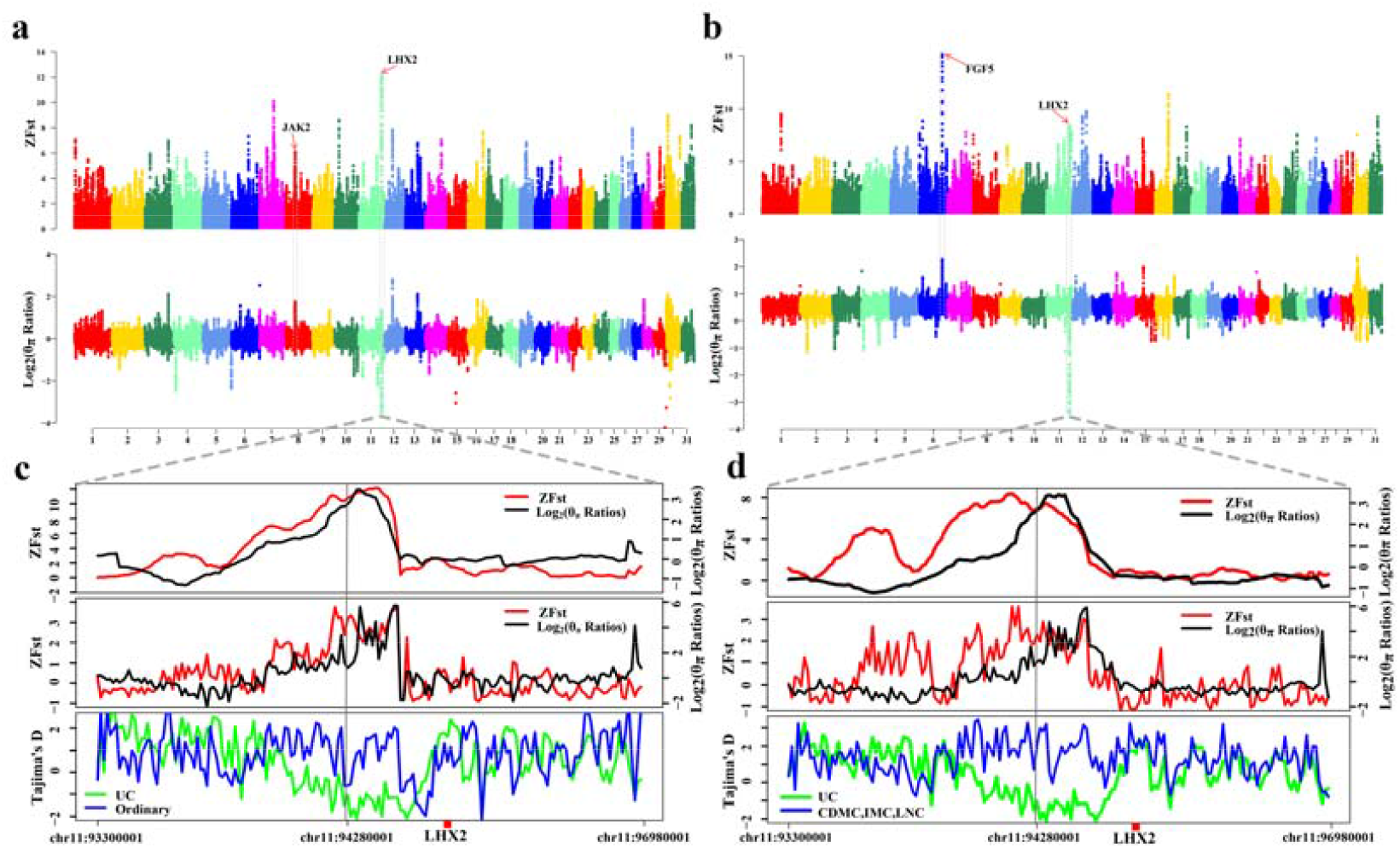
Genomic regions with selection sweep signals in domestic goats. **(a)** Manhattan plot of the genome-wide distribution of pairwise ZFst and log_2_(θ_π_ Ratios) between UC and ordinary goats (YNBB, GZB, JTB and CDB) using a 150 kb window size and a 10 kb step size. **(b)** Manhattan plot of the genome-wide distribution of pairwise ZFst and log_2_(θ_π_ Ratios) between UC and other cashmere goats (CDMC, LNC and IMC) using a 150 kb window size and a 10 kb step size. **(c)** Zooming in on the peak signal region on chromosome 11. Log_2_(θ_π_ Ratios), ZFst values, and Tajima’s D values around the *LHX2* region (Fig. 3a) using a nonoverlapping 10 kb sliding window. The black and red lines represent log_2_(θ_π_ Ratios) and ZFst values, respectively; the blue and green lines represent the ordinary goats and UC goats, respectively. All three methods indicate a strong selection signature around the *LHX2* region. **(d)** Log_2_(θ_π_ Ratios), ZFst values, and Tajima’s D values around the *LHX2* region (Fig. 3b).

In TreeMix analysis, we found that cashmere goats and ordinary goats clustered into two groups (Fig. 2b). We observed two migration edges among clusters from LNC to CDMC and from YNBB and GZB to the cashmere goat (Fig. 2b), which can be explained by the fact that CDMC goats were crossbred between goats of the Qinghai-Tibet Plateau and LNC.

Seven high-coverage samples were chosen to infer the effective population size (Ne) over the last 10^6^ years using the pairwise sequential Markovian coalescent (PSMC) method(Li and Durbin 2011). PSMC analysis suggested that the goats suffered at least two bottlenecks (approximately 7000 and 18000 years ago), which resulted in a severe reduction in the effective population size (Fig. 2c). Interestingly, the results of the PSMC analysis of goats were like the demographic history of the sheep(Yang et al. 2016).

### Genome-Wide Selective Sweeps

The results of the genetic structure analysis showed that the two cashmere goats (UC and CDMC) had apparent admixture with some other ordinary breeds. In contrast to CDMC bred through crossbreeding between Qinghai native goats and LNC, UC is originally from the Ujumuqin region and is continually selected for cashmere traits. Therefore, the distinct genetic background of UC raised the question of whether there is a different selective sweep region for the cashmere trait. We then performed a selective sweep analysis with the 12 UC and 58 ordinary goat genome sequences by estimating pairwise genetic differentiation (Fst) and nucleotide diversity differences (θπ) in 150 kb sliding windows along the genome. Using the top 1% of ZFst values and log_2_(θ_π_ Ratios) cutoffs (ZFst > 4.01, the absolute value of log_2_(θ_π_ Ratios) > 0.85), we identified 201 candidate genes associated with cashmere traits (Fig. 3a, Supplementary Table s1). We found that the most prominent ZFst signature was on chromosome 11 (Fig. 3a), spanning ∼1,100 kb region (93.6–94.7 Mb). This signature was also supported by the ZFst, log_2_(θ_π_ Ratios), and Tajima’s D statistics (Fig. 3c). However, we did not identify the selection signal harboring *FGF5*, which has been identified in previous reports(Cai et al. 2020). We next used UC and other cashmere goats to perform selective sweep analysis and identified 263 genes corresponding to selective sweeps (Fig. 3b, Supplementary Table s2). Interestingly, both selection signals on chromosome 6 and *FGF5* were simultaneously identified in these populations. Among all the candidate genes, eight (*EDA, MyD88, CD14, IL33, TNFRSF19, LHX2, AR, STK3* and *JAK2*) were located in the NF-kappaB signaling pathway and eleven (*TCF7L1, WNT8B, BTRC, AMER2, TEL2, TEL5, TEL6, LHX2, CXXC5, NOTCH1* and *VCAN*) were found in the Wnt/ß-catenin signaling pathway (Fig. 4). These signaling pathways play a central role in regulating hair follicle morphogenesis, stem cell differentiation and hair cycle(Huelsken et al. 2001; Rishikaysh et al. 2014). The strongest selective sweep localized on chromosome 11, harbored DENN domain-containing protein 1A (*DENND1A*), Crumbs cell polarity complex component 2 (*CRB2*) and LIM homeobox 2 (*LHX2*), in which *DENND1A* and *CRB2* are associated with polycystic ovary syndrome (PCOS) and maintenance of apicobasal polarity in retinal pigment epithelium, respectively, but without evidence in hair development (McAllister et al. 2014; Paniagua et al. 2021); while another gene *LHX2* near *DENND1A*, plays important role in hair follicle development.

**Fig. 4.**
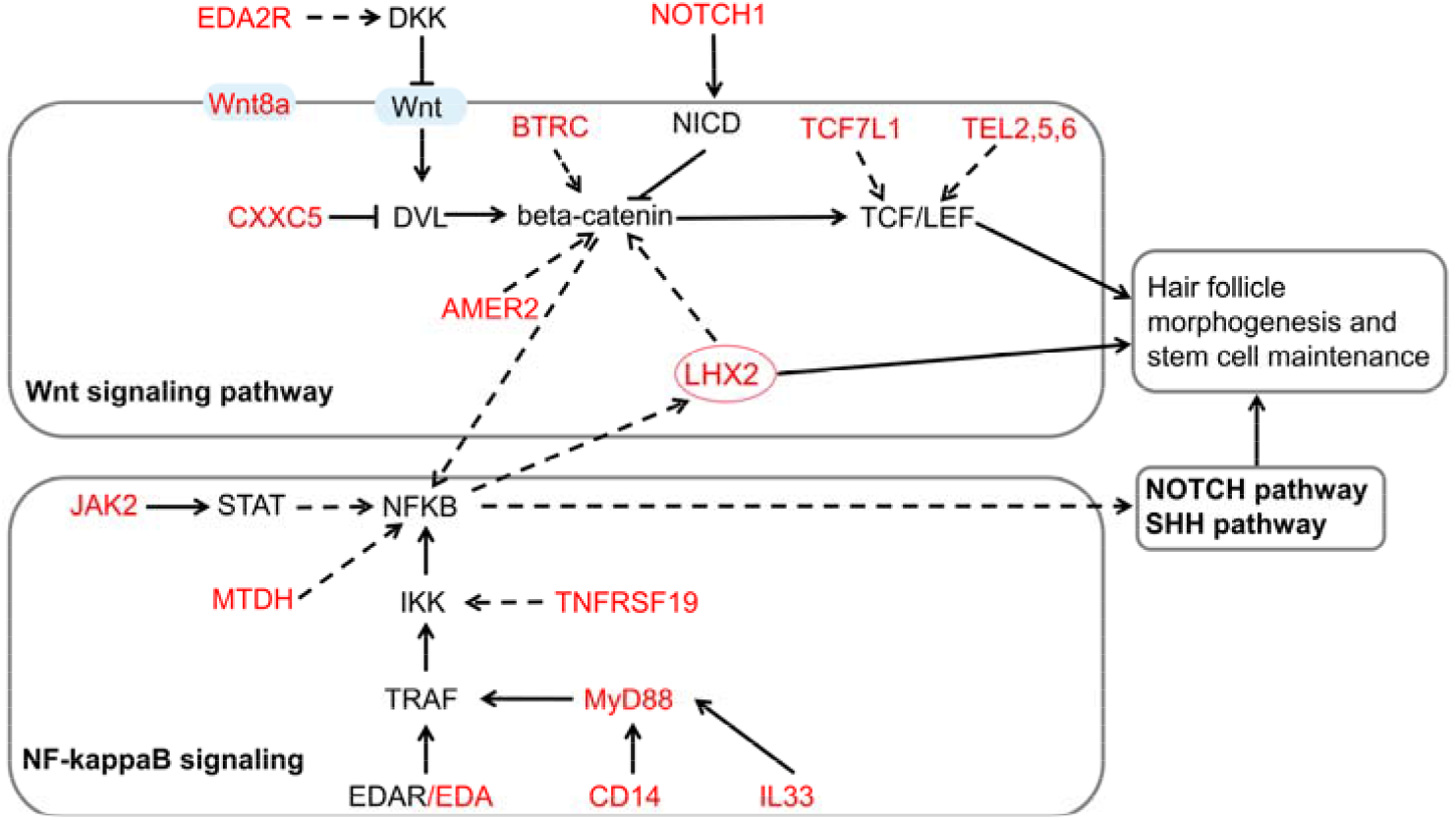
Schematic mechanisms of signaling pathways involved in cashmere fiber development. The names of the KEGG pathways are shown in bold. The candidate genes positively selected in the two methods of Fst and θπ ratio tests are shown in red. The solid block arrows represent direct effect, and the dashed black arrows indicate an indirect effect. The blunt head arrow indicates an inhibition effect.

To confirm the reliability of our findings, we further used other cashmere goats (LNC, IMC and CDMC) and ordinary goats to perform sweep selection analysis. The results showed that all previously-detected functional genes for cashmere traits including *FGF5, EDA2R* and *STIM1*, were re-identified (Supplementary Fig. 6)(Cai et al. 2020; Li et al. 2020). In addition to the cashmere trait, we also investigated the selection sweep for coat colors and confirmed previously reported genes *KIT, KITG, IRF4* and *ASIP* for coat color using different goat breeds (Supplementary Fig. 7 and Supplementary Fig. 8). Furthermore, we also identified previously reported 100 kb copy number variants compassing *KIT* (chr6:70,859,258-70,959,918) and ∼154 kb copy number variants within *ASIP* (chr13:63,226,824-63,381,501) with single base-pair resolution (Supplementary Fig. 10 and Supplementary Fig. 11).

### Plausible Causative Mutation near *LHX2*

We inspected all variants within exons to identify the potential causal mutation around the *DENND1A-LHX2* locus; however, no coding variants were found. Strikingly, a 582 bp (chr11: 94,272,264-94,272,845 bp) deletion was identified in the 13th intron of *DENND1A* and 367 kb upstream of *LHX2* with Integrative Genomics Viewer (IGV) (Fig. 5a). We tried to genotype of the deletion variant of everyone in different populations, but it is not obvious to distinguish heterozygous type from homozygous wide type due to limited read counts with IGV viewer, but it is quite confident to identify homozygous wide type. We then compared the frequencies of the homozygous deletion variant in different goat populations. The analysis results showed that the 582 bp deletion near *LHX2*(named 582del) had a higher frequency (52.9–100 %) in cashmere goats as compared 0.00%-40.00% in ordinary goats (Fig. 5b and Supplementary Table 6). We also identified a 504 bp deletion (chr6: 95,454,685-95,455,188 bp) in the *FGF5* locus (named 504del), which is consistent with previous reports (Fig. 5a) (Cai et al. 2020). However, the frequency of the 582del in the ordinary goats was much higher than that of the 504del (Fig. 5b, Supplementary Table 7). More interestingly, we found that the 582del has a high frequency in the IRWG population (80.9 %), while the 504del was absent, which is also consistent with previous research(Cai et al. 2020). Since we could not accurately distinguish the heterozygous from wild genotypes using IGV viewer, we designed PCR primers for detecting the heterozygosity of the two deletions in several goat populations. It showed that the frequency of homozygosity was consistent with that by IGV viewer (Fig. 5c, Fig. 5d, Supplementary Table 8, and Supplementary Table 9), but some breeds had a higher frequency of heterozygosity, such as in UC (504del+/-: 47.5 %) and JNG goats (504del+/-: 66.7 %; 582del+/-: 66.7 %). Finally, we detected the 582del in several ancient goat samples (7/21) and ibex samples (1/3) (Supplementary Fig. 13 and Supplementary Fig. 14), indicating that the 582del is an older mutation that occurred much earlier than the 504del. In addition, we compared the genotypes of the target genomic region for cashmere and ordinary goats. The result showed that the genotype patterns of cashmere goats were highly like that of wild goats, but different from that of ordinary goats (Supplementary Fig. 15).

**Fig. 5.**
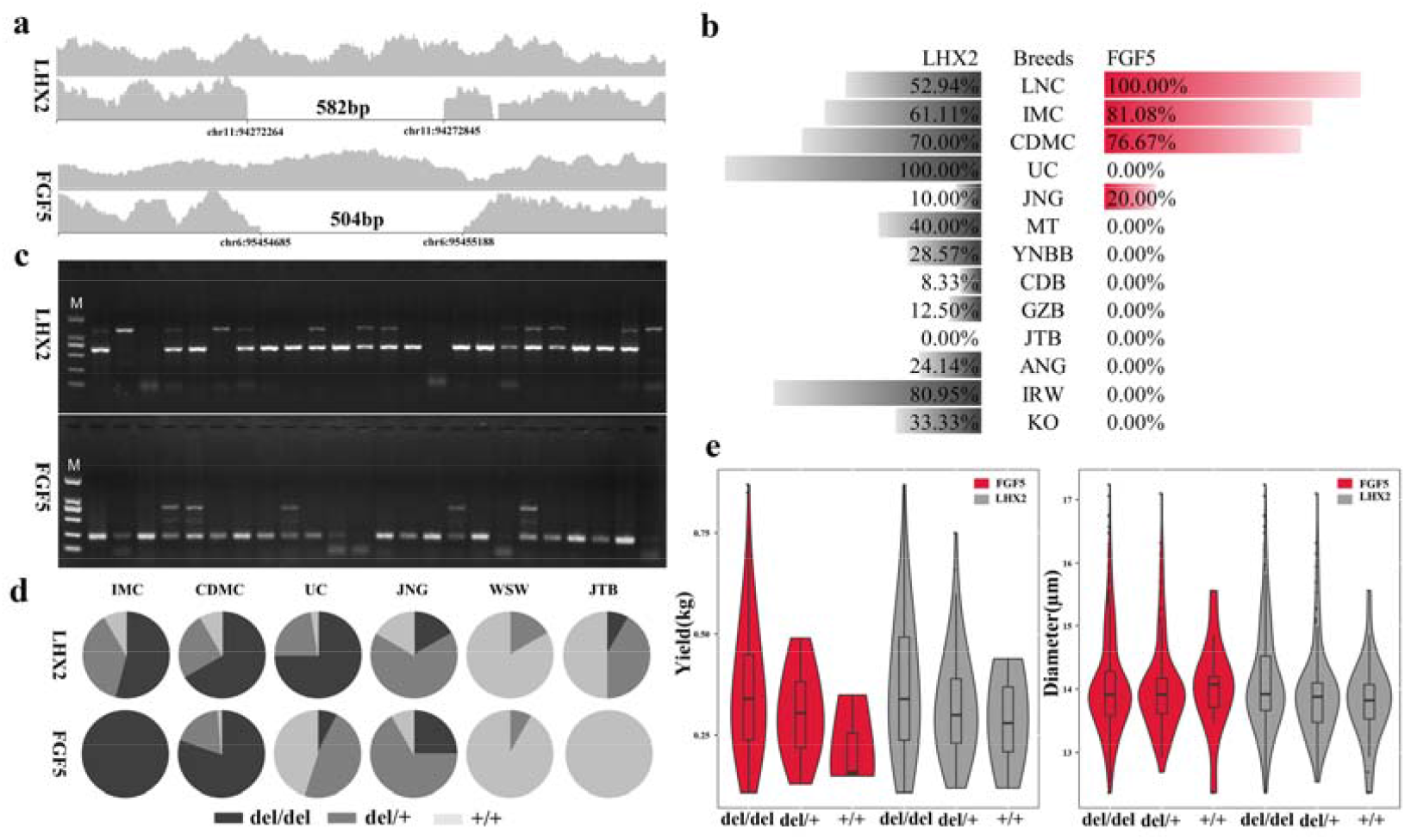
The *LHX2* and *FGF5* deletions in different goat breeds and their association with cashmere traits in CDMC. **(a)** The identified 582 bp deletion near *LHX2* and the previously reported 504 bp deletion near *FGF5* based on read coverage. **(b)** Homozygous genotype frequency of the two deletions determined by IGV viewer **(c)** PCR amplification of the two deletion variants. **(d)** Distribution of the 582 bp deletion near *LHX2* and the 504 bp deletion of *FGF5* genotypes. del/del, deletion/deletion; del/+, deletion/wild type; +/+, wild type/wild type. WSW (Wushan White goat from southwest China). **(e)** The association of the 582 bp deletion near *LHX2* and 504 bp deletion of *FGF5* with cashmere yield and diameter in the CDMC goat population.

To further evaluate whether these two deletion variants were related to cashmere traits, we selected 235 CDMC goatswith cashmere yield (Supplementary Fig. 22, Supplementary Table s8) and 581 CDMC goats with fiber diameter records (Supplementary Fig. 23, Supplementary Table s9) for association analysis. The association results showed that the 582del and the 504del was significantly associated with cashmere production (*P* value = 0.0061 and *P* value = 0.0113 for the 582del and the 504del, respectively). The 582del was also significantly associated with fiber diameter (*P* value = 2.74e-04), but the 504del was not (Fig. 5e, Supplementary Table s8, and Supplementary Table s9). Interestingly, the interactions effect of the 582del and 504del were significantly associated with fiber diameter (*P* value = 6.1e-06) (Supplementary Table s9), suggesting crosstalk between the two genes.

### Biological Function of the 582 bp Deletion

Analysis of the 582del deletion region using the BLAST program revealed that it is not a highly conserved element but was found in the genomes of primate and ungulate species (Fig. 6a). By checking the UCSC Genome Browser (https://genome.ucsc.edu/), we found many cis-regulatory elements and H3K27Ac marks (often enriched in enhancer regions) upstream of this deleted region (Fig. 6b). Interestingly, we identified a CTCF-binding site and two distal enhancer-like signatures, which were close to the 582bp deletion sequence (Supplementary sequence 1). Using previously reported RNA-seq data(Wu et al. 2020), we found that the expression of *LHX2* in the different fetal stages of cashmere goats exhibited an up-regulation pattern during hair follicle development (Fig. 6c, Supplementary Fig. 16). Furthermore, some functional enhancers of *LHX2* were also identified in the genome regulatory blocks of the *CRB2-LHX2* loci(LeeBrenner and Venkatesh 2011). This evidence suggests that the deletion sequence may function as an insulator to block the *LHX2* enhancer function.

**Fig. 6.**
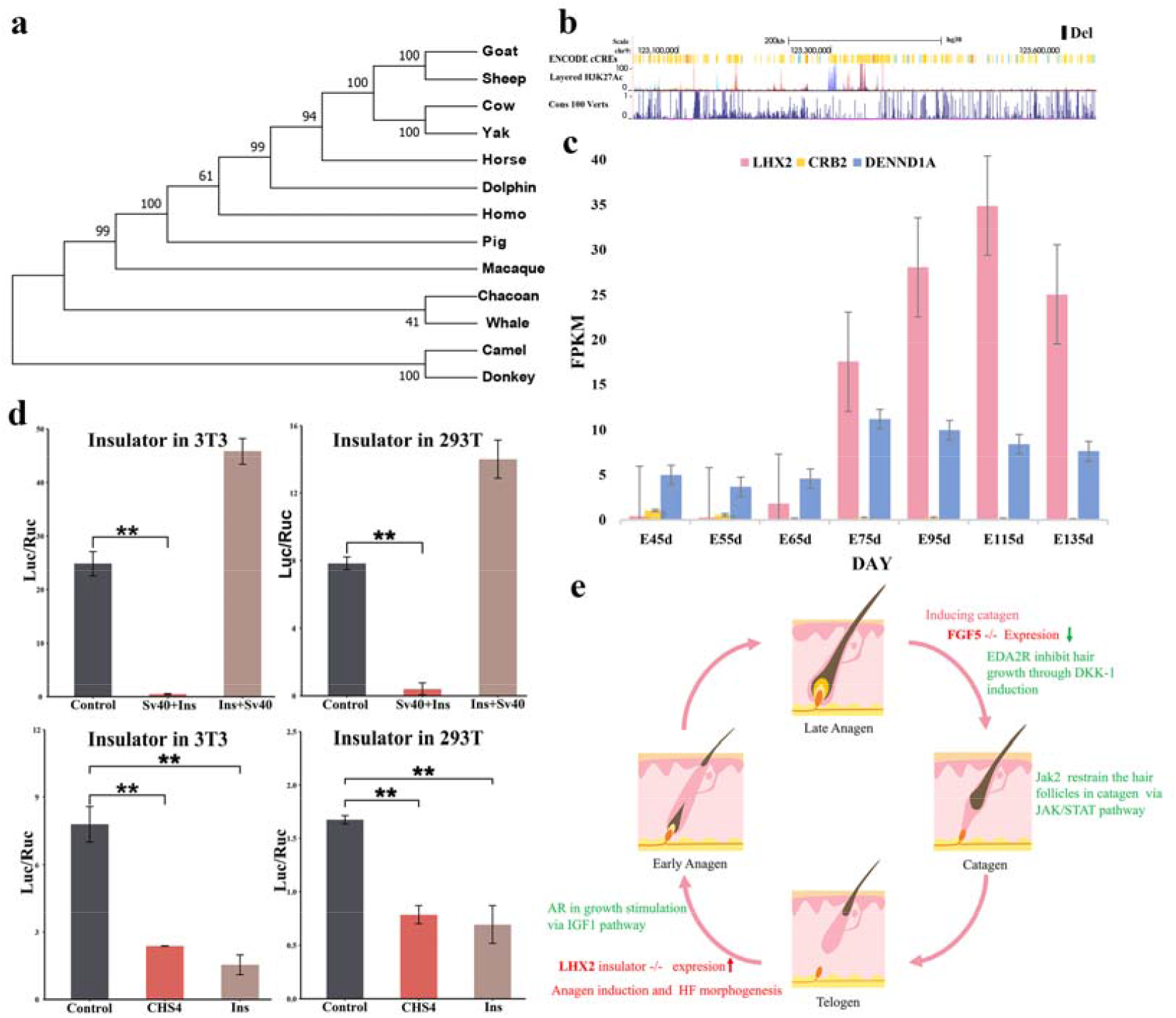
Effects of *LHX2* deletion sequence on reporter gene expression in mouse NIH3T3 cell line and human 293T cell line. **(a)** Phylogenetic tree of 13 primates and ungulates for the detection of the 582 bp deleted sequence. **(b)** Regulatory elements, epigenomic signals, and conservation scores on selective sweep region in chromosome 11. The black box represents the deletion variant location of *LHX2*. **(c)** The expression of *LHX2, CRB2* and *DENND1A* in different stages of prenatal skin in cashmere goats. Boxes of different colors are used to represent genes. The expression data is downloaded from a previous report(Wu et al. 2020). **(d)** A dual-luciferase assay using NIH3T3 cell and 293T cell shows the *LHX2* upstream deletion sequence blocks the activity of luciferase. Data are shown as the mean ± standard error. The P-value was calculated using Student’s *t*-test. **(e)** The molecular mechanisms of cashmere fiber development. In detail, the deletion of the *LHX2* insulator increases the expression of *LHX2* and promotes the cashmere fiber growth at the anagen stage, and *AR* could participate in hair growth at the anagen stage through the *IGF1* pathway(Grymowicz et al. 2020), while the deletion of the 504 bp enhancer reduces the expression of *FGF5*, thereby inhibiting growth regression; *JAK2* and *EDA2R* may also participate in this stage via the JAK/STAT and Wnt pathways, respectively.

To confirm the insulator function of this sequence, we synthesized a 551 bp DNA fragment (named Ins, Supplementary sequence 2) and subsequently inserted it downstream and upstream of the SV40 promoter in the pGL3 plasmid (Supplementary Fig. 17). The Ins vectors with the Renilla luciferase vector, phRL-TK, were transiently co-transfected into human 293T cells and mouse 3T3 cells. After 48 h, the firefly and Renilla luciferase activities of the lysate were measured, and the ratio of firefly luciferase activity to Renilla luciferase activity was calculated for each sample. Our data showed that the Ins fragment decreased the expression of firefly luciferase by more than 90 % in the downstream group of both cell types and increased expression in the upstream group (Fig. 6d). In addition, we inserted the Ins downstream of the SV40 enhancer in the pGL3 plasmid (Supplementary Fig. 17) and chose the well-known insulator cSH4 as a control to quantify the efficiency of the Ins. Our data showed that the Ins fragment decreased the expression of firefly luciferase by approximately 50 % in human 293T cells, which was comparable to that of cSH4 (Fig. 6d). In addition, a similar result of the Ins activity was validated in the mouse 3T3 cells but decreased more sharply than the cSH4 group (Fig. 6d). These results suggested that the 551 bp DNA fragment played an enhancer-blocking function in mammary cells.

To describe the function mode of these selective sweep genes on the development cycle of cashmere, we downloaded and analyzed the transcriptome data of skin tissue during the prenatal stages and one-year cashmere cycle. The transcriptome data showed that *AR* and *EDA2R* were not expressed in embryonic cashmere goat skin, and *JAK2* gene expression was opposite to that of *LHX2* (data from Wu(Wu et al. 2020), Supplementary Fig. 18). Furthermore, *JAK2, EDA2R* and *FGF5* exhibited similar expression patterns in the skin transcriptome data of different months of the one-year cycle, while *LHX2* and *AR* had similar expression patterns (PRJNA470971) (Supplementary Fig. 18 and 19). It is known that cashmere fiber growth has three different periods in one year: a growth period (March–September), a regression period (September–December), and a resting period (December–March)(Yang et al. 2020). Therefore, the deletion of the *LHX2* insulator increases the expression of *LHX2* and promotes cashmere fiber growth at the anagen stage, while deletion of the *FGF5* enhancer reduces the expression of *FGF5*, inhibiting the regression. The other candidate genes may affect cashmere fiber growth through uninvestigated regulation modes. Overall, we propose a possible molecular model for cashmere fiber formation: *LHX2* and *AR* may be involved in maintaining hair growth at the anagen stage, whereas *JAK2, EDA2R* and *FGF5* function in the destructive phase (catagen) through the highest expression in September (Fig. 6e).

## Discussion

In this study, we resequenced the genomes of 42 cashmere goats and 78 ordinary goats. Population analyses revealed that goats in southwest China are different in the genome from goats in West Asia and North China, which is consistent with a recent study(Cai et al. 2020). In the present study, we observed a genetic introgression from the LNC breed into the CDMC breed, which was confirmed by the origin and breed practice of CDMC (using Chaidamu goats as female parents, and LNC as male parents to breed new cashmere goats, named CDMC)(Zheng-luZhaeng-kui and DARIQIBU 2003). In addition, the introgression from the Yunnan-Kweichow Plateau into the cashmere goat breeds suggests that goats from the Qinghai-Tibet Plateau and Mongolian Plateau, as well as from the Yunnan-Kweichow Plateau, may have a common ancestor or have a genetic admixture in their early years. Similar introgression results have also been reported in sheep populations from these regions(Yang et al. 2016). In contrast, Angora goats have a signature of genetic admixture with some Chinese goats (Fig. 2a), which has also been reported previously(Ryder 1993). In addition, although KO has large genome differences with other goat breeds(Kim et al. 2019), a clear signature of genetic admixture between KO and JT, JNG, and MT was observed, which may be related to the historical human migration between China and Korea. Finally, we observed CDMC and IMC goats presenting a genetic admixture of UC and LNC. LNC was bred in the 1980s from six counties in the eastern mountainous area of Liaoning province in China, famous for its high cashmere yield. UC was originally bred from Ujumuqin white goat in the region of Ujumuqin grassland for selecting the cashmere trait in 1994. Unlike LNC, UC has a closer genetic distance to goats in the middle region of China (Fig. 2a). These results indicate that UC may have a more unique genetic background than other cashmere goats.

By performing a whole-genome selection scan, we discovered a novel selective sweep region on chromosome 11 that appears to have undergone extremely strong selection in UC. This novel selective sweep region is located in a conserved linkage block, containing three genes: *DENND1A, CRB2* and *LHX2*(LeeBrenner and Venkatesh 2011). Among these genes, *LHX2* functions as a transcriptional activator in hair follicle stem cells and is an essential positive regulator of hair formation(RheePolak and Fuchs 2006; Tornqvist et al. 2010). Furthermore, the expression of *LHX2* was upregulated during hair follicle differentiation, and the expression abundance was constant throughout the development cycle in secondary hair follicles of cashmere goats(Wu et al. 2020; Yang et al. 2020). In contrast, *DENND1A* and *CRB2*, which function in endocytic trafficking to mediate the recycling of selective cargos and early embryonic development, respectively(Xiao et al. 2011; Shi et al. 2019), were expressed lowly in the fetal stage of goat skin and showed very low expression during the cycle of adult cashmere fiber growth (Fig. 6c). These results indicate that *LHX2* is a functional gene in this selective sweep region.

We then reported an upstream deletion of *LHX2*, which carries a potential cis-regulatory insulator region (Fig. 6d). In contrast to the deletion allele of *FGF5*(Cai et al. 2020), the deletion allele of *LHX2* was found in all goats investigated, including wild and domesticated goats, but allele frequency was higher in northern and wild goats than in goats in other areas (Fig. 5b and Fig. 5d). Notably, this deletion was also found in the ancient goat and ibex samples (Supplementary Figs. 13 and 14). Ibex goats may also have dense underwool, like cashmere goats(Ryder 1993). These results indicate that the deletion allele of *LHX2* occurred in the pre-domestication stage of goats and might be traced back to the early stage of goat speciation to cold climate adaptation. Further, we did not detect the 582 bp deletion in the same region of sheep, but we cannot rule out the possibility that we tested too few sheep (Supplementary Fig. 20). In contrast, the deletion allele of *FGF5* is most likely to occur during the domestication stage of goats to obtain higher cashmere production.

Overall, we described a possible model for the origin and diffusion of cashmere-related mutations during the selection evolution of cashmere goats, helping us understand the molecular basis of cashmere trait formation (Fig. 7).

**Fig. 7.**
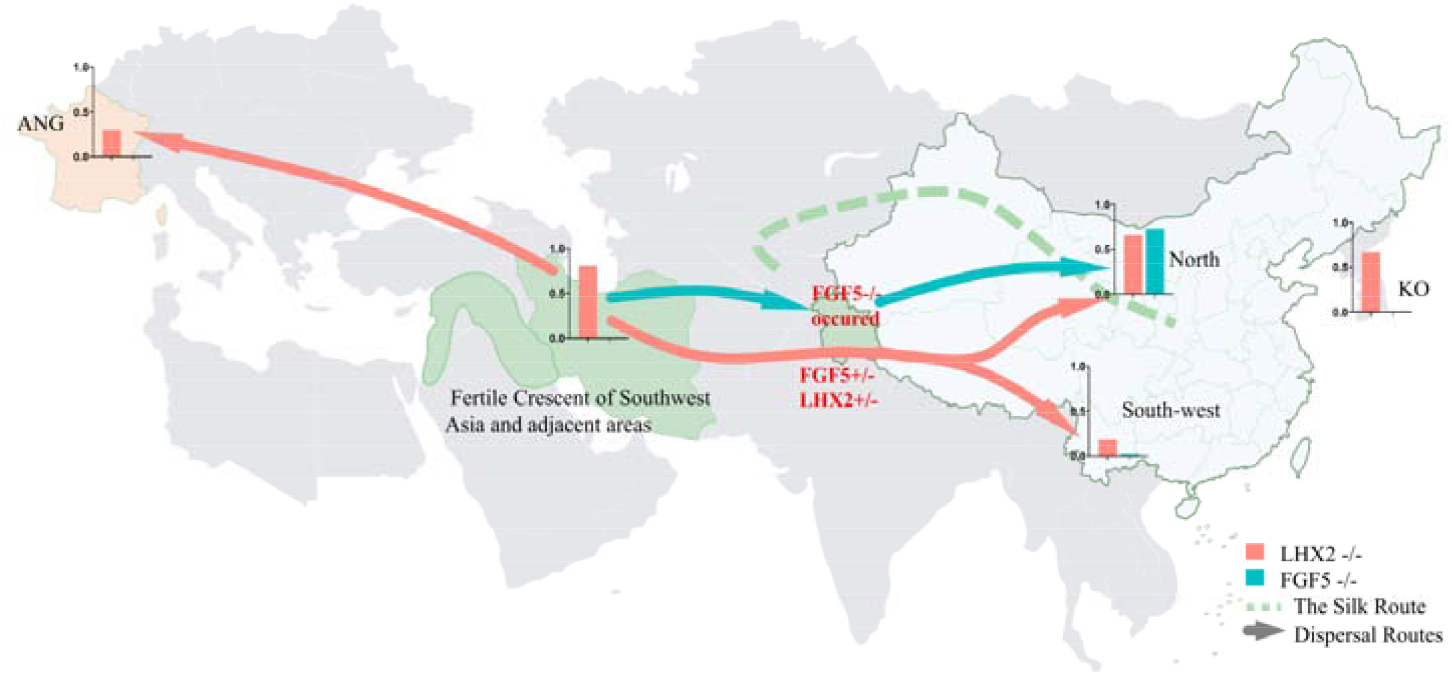
Origin and diffusion of causative mutations for *LHX2* and *FGF5* in wild and domesticated goats. Some wild goats acquired the deletion allele of *LHX2* at a very early stage. Because this deletion is beneficial to the development of cashmere to adapt to the cold environment, goats obtained higher genotype frequencies in the north than in the south. The *FGF5* enhancer deletion may have occurred during artificial selection or secondary domestication in the Kashmir region of Afghanistan and gradually migrated north of the Mongolian Plateau. This enhancer deletion was fixed during the long-term artificial selection. The low frequency of the *FGF5* enhancer deletion genotypes in the southwestern region may be related to the genetic admixture between the Qinghai-Tibet Plateau and the Yunnan-Kweichow Plateau goats.

Multiple sequence alignment revealed that the selective sweep region contains many CTCF binding sites and other cis-regulatory elements, as well as the H3K27ac epigenetic mark, located upstream of the 582bp deletion. CTCF is a highly conserved zinc finger protein, required for insulator function in mammals(WestGaszner and Felsenfeld 2002), while H3K27ac is a marker for active enhancers and a great indicator of enhancer activity(Creyghton et al. 2010), suggesting that the selective sweep region contains some hair-related transcription regulatory elements. Our biological function experiments showed that the 582bp deletion region contains an insulator for blocking the upstream enhancer role with higher efficiency in skin-related cells (NIH3T3 cells) than cSH4. An insulator is a long-range regulatory element, which protect an expressing gene from its surroundings through two ways: blocking the action of a distal enhancer on a promoter and acting as “barriers”that prevent the advance of nearby condensed chromatin (WestGaszner and Felsenfeld 2002). Previous report described that four of the eight conserved noncoding elements (CNEs), approximately 222 kb to 619 kb upstream of human *LHX2*, functioned as tissue-specific enhancers in specific regions of the central nervous system and the dorsal root ganglia (DRG), recapitulating partial and overlapping expression patterns of *LHX2* and *CRB2* genes (LeeBrenner and Venkatesh 2011). In our research, we found that the insulator is situated approximately 367 kb upstream of goat *LHX2* in the 13th intron of *DENND1A*. Therefore, we speculate that this insulator can regulate the expression of the *LHX2* gene by blocking the upstream enhancers, and the deletion of the insulator can increase the expression of the *LHX2* gene, thus increasing cashmere production. Subsequently, association analysis confirmed the significant relationship between the deletion genotype and cashmere traits, including cashmere production and fiber diameter (Fig. 5e). The interaction effect of the two deletion variants significantly affected the diameter of the cashmere fiber, indicating that the two genes have a synergistic effect on cashmere fiber development.

In addition to *LHX2* and *FGF5*, we also identified a few positively selected genes related to cashmere traits, four genes (*AR, JAK2, EDA2R* and *STK3*) affecting I-kappaB kinase/NF-kappaB signaling, and five genes (*NOTCH1, TCF7L1, AR, VCAN* and *WNT8B*) influencing the canonical WNT signaling pathway (Supplementary Table s7). We found that *NOTCH1, TCF7L1* and *STK3* exhibited similar seasonal expression patterns with *JAK2, FGF5* and *EDA2R* in the skin, but *VCAN* and *WNT8B* genes showed low expression (Supplementary Fig. 21; PRJNA470971). NF-kappaB and WNT signaling pathways play a central role in hair follicle development and regeneration(KishimotoBurgeson and Morgan 2000; Krieger et al. 2018; Choi 2020). Interestingly, *LHX2* is also regulated by NF-κB signaling to promote primary HF morphogenesis(Tomann et al. 2016). *EDA2R* is a divergent gene between cashmere goats (excluding UC goats) and ordinary goats, consistent with previous research(Cai et al. 2020). *EDA2R* is highly expressed in the late anagen phase and may inhibit hair growth by inducing *DKK-1* expression(Kwack et al. 2020). Furthermore, the *AR* gene and *JAK2* are novel genes identified in cashmere traits.

*JAK2* can restrain the hair follicles in catagen via the JAK/STAT pathway with up-regulation in catagen and telogen stages(Harel et al. 2015; Tao et al. 2020). In contrast, AR has paradoxically different effects on hair follicles(Randall 2008). In cashmere goats, *AR* is expressed in the early anagen stage and may stimulate cashmere growth via the *IGF-1* pathway(Inui and Itami 2013). These positively selected genes may be involved in cashmere trait formation through the model described in Fig. 6e.

## Conclusions

Through population genomics and selective sweep analyses of cashmere and ordinary goats, we identified a novel causative mutation upstream of *LHX2* that functions as an insulator to block the enhancer of *LHX2*. In contrast to enhancer deletion of *FGF5*, this insulator deletion also retained high allele frequency in wild goats, ancient goats, and ibex. We also found that the insulator deletion was associated with two cashmere traits in the CDMC goat population. The positively selected genes were enriched in the NF-κB and Wnt pathways, which play a central role in hair follicle development. This study not only provides the first evidence of how cashmere fiber growth could be regulated during the evolution of goats but also offers valuable molecular markers for the genetic improvement of cashmere traits in goats.

## MATERIALS AND METHODS

### Samples Collection

We collected 120 domestic goats (C. hircus), representing eight geographically diverse breeds in China, which include 30 CDMC from Chaidamu City of Qinghai Province; 11 UC from Ujimqin Banner of Inner Mongolia Province; 8 GZB from Guiyang City of Guizhou Province; 11 YNBB from Lanping County of Yunnan Province; 10 MT from Shangqiu City of Henan Province; 10 JNG from Jining City of Shandong Province; 10 JTB from Jintang County of Sichuan Province; 12 CDB from Chengdu City of Sichuan Province.. Details of the samples used in this study are givened in Supplementary Table 1. All animal procedures were approved by the life ethics and biological safety review committee of BGI (NO. FT 18041), and were carried out in accordance with the approved guidelines.

### DNA Extraction and Sequencing

The ear or blood tissues of CDMC, UC, JT, GZB, JNG, YNBB, CDB and MT were collected on-site and stored in an alcohol sampling tube or blood collection tube. A tissue DNA extraction kit was used to extract the genomic DNA from the samples, electrophoresis was used for integrity detection, and Qubit was used for concentration determination. Library construction for resequencing was performed with 1-3 µg of genomic DNA using standard library preparation protocols and insert sizes from 200–400 base pairs (bp). All 120 goats were sequenced on the BGISEQ-500 platforms with PE100 (CDMC, UC, JT, YNBB, CDB, MT) and on the Illumina HiSeq2500 platforms with PE150 (GZB, JNG). Besides, we downloaded published genomic data of modern, wild and ancient goats form the NCBI deposit (Supplementary Table 2, Supplementary Table 1).

### Read Alignment and Variant Calling

We obtained raw data from a total of 236 goat samples, including 120 sequenced and 116 downloaded. All raw data were first filtered and trimmed using SoapNuke (version 1.5.0) (https://github.com/BGI-flexlab/SOAPnuke) (Chen et al. 2018), if any of the following criteria were met: (1) reads containing adapter and poly-N; (2) reads whose low-quality base ratio (base quality less than or equal to 5) is more than 50%; (3) reads whose unknown base (‘N’ base) ratio is more than 10%. Clean reads from all individuals were aligned to the goat reference genome ARS1 (https://www.ncbi.nlm.nih.gov/assembly/GCF_001704415.1/)(Hassanin" www.ncbi.nlm.nih.gov/assembly/GCF_001704415.1/)(Hassanin et al. 2010) by the Burrows-Wheeler Aligner (BWA version 0.7.12). SAMtools(Li et al. 2009) was used to convert the file format from SAM to BAM and filter the unmapped and non-unique reads. Picard (version 1.54, http://broadinstitute.github.io/picard/) was used to sort the BAM files and remove potential PCR duplications if multiple read pairs had identical external coordinates.

GATK (version 4.0.11, https://github.com/broadinstitute/gatk) tools were used in the whole process of variant calling. After mapping, the “HaplotypeCaller”, “CombineGVCFs” and “GenotypeGVCFs” in GATK4 were used to detect SNPs and Indels with default parameters. The output VCF File was then screened for SNPs using “SelectVariants function” of GATK4. SNPs and Indels were separated using the GATK tool “SelectVariants” and subjected to rigorous processing to exclude false positives. To obtain high-quality SNPs, we carried out SNP filtering at two stages. SNPs exclusion criteria(Choi et al. 2015) were as follows: (1) hard filtration with parameter “QD < 2.0 || ReadPosRankSum < −8.0 || FS > 60.0 || MQ < 40.0 || SOR > 3.0 || MQRankSum < −12.5 || QUAL < 30;” (2) “--max-missing 0.9 --maf 0.05 --min-alleles 2 --max-alleles 2.” Finally, ∼ 13 million high-quality SNPs were remained for further analysis.

### Population Structure Analysis

We used 12,861,877 high-quality SNPs (autosome) to PCA analysis. PLINK (version 1.9)(Purcell et al. 2007) was used to calculate the principal components, using function “--vcf vcf --out pca –pca --chr-set 29 --allow-extra-chr”.

An individual-based NJ tree was constructed for the 236 goats based on the p-distance, with one outgroup (IRW, Iranian wild goat) using the software VCF2Dis (version 1.09, https://github.com/BGI-shenzhen/VCF2Dis) and PHYLIP (version 3.69) (http://evolution.genetics.washington.edu/phylip.html). The Linux command line of converting to a matrix is “VCF2Dis -InPut vcf -OutPut p_dis.mat”; the Linux command line of constructing a phylogenetic tree is “ fneighbor -datafile p_dis.mat -outfile goat.tree.txt -matrixtype s -treetype n -outtreefile goat.n.tree.tree”.

Population structure was analyzed using the ADMIXTURE (version 1.23)(AlexanderNovembre and Lange 2009; Alexander and Lange 2011) program which implement a block-relaxation algorithm. To explore the convergence of individuals, we predefined the number of genetic clusters K from 3–7 and ran with cross-validation error (CV) procedure. Default methods and settings were used in Admixture analysis. The analysis process includes the following three steps:

(1) Converting VCF format to PLINK format. The Linux command line was “vcftools

--vcf vcf --plink --out goat.plink”;

(2) Using PLINK for further filtering with Linux command line “plink --noweb --file plink --geno 0.05 --maf 0.05 --hwe 0.0001 --chr-set 29 --make-bed --out QC”; after this step we get the corresponding bed file.

(3) ADMIXTURE v1.23 for population structure analysis with “admixture --cv QC.bed $k | tee log${k}.out”; then the results were plotted using R.

PopLDdecay (https://github.com/BGI-shenzhen/PopLDdecay)(Zhang et al. 2019) for linkage disequilibrium decay analysis was based on variant call format files.

### Demographic History

A population-level admixture analysis was conducted in the TreeMix (version 1.12)(Pickrell and Pritchard 2012). The program inferred the ML tree for 12 goat breeds (215 individuals) and an outgroup (Bezoars,21 individuals), then the residuals matrix was used to identify pairs of populations that showed poor fits in the ML tree. These populations were regarded as candidates around which we added potential migration edges, and new arrangements of the ML tree accounting for migration events were generated (Pickrell and Pritchard 2012). From one to 20 migration events were gradually added to the ML tree until 98% of the variance between the breeds could be explained. The command was ‘-i input -bootstrap -k 500 -root IRW -o output’.

We used the PSMC (version 0.6.5) (Li and Durbin 2011) method to estimate changes in the effective population size (Ne) of goat over the last one million years. The PSMC analysis was implemented in the seven samples sequenced at a high read depth (12.6∼26.94×) (Supplementary Table 11).The parameters of PSMC were set to -N25 -t15 -r5 -p “4+25^*^2+4+6”, where the parameters of generation time and mutation rate were set to 2.0 and 2.5e-8, respectively.

### Selective Sweep Analysis

In the selection sweep, we calculated the genome-wide distribution of Fst values and θ_π_ Ratios (Danecek et al. 2011) (i.e., θ_π-UC_/θ_π-Ordinary_, θ_π-UC_/θ_π-CDMC.IMC.LNC_, θ_π-LNC.IMC_/θ_π-Ordinary_, θ_π-White_/θ_π-Brown_ and θ_π-White_/θ_π-Black_) for five control group pairs, which included the UC group versus the ordinary group (GZB, YNBB, JTB, CDB), the UC group versus other cashmere (CDMC, IMC, LNC), the other cashmere group (CDMC, IMC, LNC) versus the ordinary group (GZB, YNBB, JTB, CDB), the white coat color(UC,CDMC, IMC, LNC) versus the brown coat color(CDB), the white coat color(IMC, LNC) versus the black coat color(YNBB, GZB) using a sliding-window approach (150-kb windows with 10-kb increments). The Fst values were Z-transformed, and the θπ Ratios were log_2_-transformed. We considered the windows with the top 1% values for the ZFst and log_2_(θ_π_ Ratios) simultaneously as the candidate outliers under strong selective sweeps. All of the outlier windows were assigned to corresponding SNPs and candidate genes. The signals were further confirmed by ZFst, Log_2_(θ_π_ Ratios), and Tajima’s D with a 10 kb sliding window.

### Functional Enrichment Analyses

Candidate genes under selection were defined as those overlapping sweep regions or within 500 kb of the signals. The biological function of genes within candidate regions was annotated by analyzing Gene Ontology (GO) and Kyoto Encyclopedia of Genes and Genomes (KEGG) pathways using Metascape(http://metascape.org/). Benjamini–Hochberg FDR (false discovery rate) was used for correcting the P values. The Gene Ontology categories “Molecular Function,” “Biological Process” and “Cellular Component”, and Human Phenotype (HP) categories were used in these analyses.

### Genotypic and phenotypic analysis

### Genotyping

Genotypic and phenotypic analysis DNA was extracted from the ear-tissue samples. The 582 bp deletion near *LHX2* (582del) and the 504 bp deletion of *FGF5* (504del) were genotyped by PCR amplification using the reaction condition of the 2-min pre-denaturation at 95 °C, 15s-denaturation at 95°C, 30s-annealing at 56°C, 45s-extension at 72 °C for 30 cycle, and 5-min extension at 72°C (the primers of 582 bp deletion: 5’-CAGTACGGAGCAAGTAAACGG-3’ and 5’-ACCATTCCACTTGTCCACCT-3’, the primers of 504 bp deletion: 5’-ACAGCGTGTGATCTTTTCTCTG-3’ and 5’-TCTTGGTCTGGCTGTGATCA-3’) and visualized on 2% agarose gels. The 582del and 540del were successfully genotyped in the extended population of 840 goats, including 752 CDMC, 12 IMC, 40 UC, 12 JNG, 12 Wushan white goats (from Chongqing in the southwest of China) and 12 JTB individuals.

### Phenotypic analysis

All 581 cashmere samples were collected from the lateral body part of CDMC goats, and the fiber diameter was measured by an optical-based fiber diameter analyzer (OFDA2000). Sample preparation is the most critical part of obtaining accurate results from OFDA. Briefly, the cashmere fibers are roughly aligned in parallel and cut into 2 mm by the OFDA bench guillotine, and then washed the fibers to remove grease. After drying, use a spreader to evenly spread 10-15 mg of fibers on the glass slide. Place the slide on the stage, then enlarge and scan the snippets through the optical transmission microscope. The camera system collects the fiber image, and then the fiber in it is automatically identified through image analysis technology and the diameter is measured. In addition, we measured cashmere yield from a subset of 351 CDMC goats in May 2020. The cashmere production is measured by hand-combing cashmere and weighing. First, use a thin comb to comb the feces and other dirt along the direction of the hair, then use a density-comb to comb the direction of the hair repeatedly and finally comb the hair in the reverse direction until the fallen fibers are combed. The collected cashmere is weighed with an electronic scale (range 0.3g-5kg, accuracy 0.1g). Then, we tested if the identified deletion variants were associated with cashmere diameter and production, respectively.

Totally, 581 individuals were recorded, the linear model was expressed as, y=b0+bx+sex+batch+e, where, y was the diameter; x was the deletion coding with 0, 1 and 2 representing 0, 1 and 2 copies of deletion sequence, respectively; b was the effect of deletion variant; sex was the sex effect; batch was the recorder effects, where 2 recorders were participated in the phenotyping; e was the residual error, assumed to follow normal distribution. When performed the association between the deletion variants and the production, 235 individuals were included, all were females, the linear model for the association was expressed as y=b0+bx +e. The association studies were performed with lm function of R package.

### RNAseq

RNAseq data were downloaded from the NCBI database project id SRP145408. Raw data were filtered by SOAPnuke (version 1.5.6), read base N more than 5%, and read base quality less than 20 was removed. Clean reads from the 12 individual samples were aligned to the reference genome (GCA_001704415.1 ARS1) using hisat2 (version 2.1.0). Picard-tools-1.105 was used to sort BAM files. Based on the mapped reads and goat reference genome annotation in GFF (GCF_001704415.1), StringTie (version,1.3.3b) was used to calculate the FPKM(Pertea et al. 2016).

### Dual-Luciferase Reporter Analysis

*Plasmid Constructs*, a 563 bp DNA fragment containing “5’ end - KpnI/HindIII site-Ins sequence (551bp)-XhoI/NcoI site was synthesized by TsingKe Ltd, which was subsequently cloned downstream and upstream of the SV40 promoter in the pGL3 plasmid, named as Insulator1 and Insulator2, respectively. A 796 bp DNA fragment containing “5’ end - BamHI site-Ins sequence(551bp)-SalI site-sv40 enhancer sequence-PciI site – 3’ end” (Supplementary Fig. 17) was synthesized by TsingKe Ltd, which was subsequently cloned downstream of Luc+ Poly (A) in the pGL3 promoter plasmid (Promega) and named as Ens-E. Similarly, the well-known insulator cHS4 linked to the sv40 enhancer (Supplementary Fig. 17) was cloned into a pGL3 promoter plasmid and named cHS4-E.

Cell Culture, Transfection, and Luciferase Reporter Assay. The 3T3 cells and 293T cells were maintained in Dulbecco’s modified Eagle’s medium (Gibco, Waltham, MA, USA) with penicillin (100 units/ml), streptomycin (100 µg/ml), and 10 % fetal bovine serum at 37 °C in a humidified atmosphere of 95 % air and 5 % CO_2_. Before transfection, 3T3 and 293T cells were seeded into 24-well plates at 1.0 × 104 cells/well. After overnight attachment, transfections were performed using Lipofectamine 2000 (Invitrogen, CA, USA) according to the manufacturer’s instructions. In this experiment, the two types of cells were divided into eight groups, respectively. For each group, plasmid of pGL3-control (Promega), Insulator1 and Insulator2 or Ins-E and cHS4-E were co-transfected with phRL-TK (Promega) at a ratio of 10:1 (0.8 ug: 0.08 ug) into the 3T3 and 293T cells, respectively. The cells were harvested two days after co-transfection, and luciferase activity was evaluated using the Dual-Luciferase Reporter Assay System (Promega, #E1910). Firefly luciferase data were normalized to Renilla luciferase activity. For this assay, pGL3-control was used as a positive control and Lipofectamine 2000 was used as a blank control. The cHS4 insulator was chosen as a reference to quantify the efficiency of the insulator function.

## Data availability

The data that support the findings of this study have been deposited into CNGB Sequence Archive (CNSA)(Guo et al. 2020) of China National GeneBank DataBase (CNGBdb)(Chen et al. 2020) with accession number CNP0001896

(https://db.cngb.org/cnsa/project/CNP0001896/reviewlink/).

## Acknowledgements

This work was supported by the Science and Technology Innovation Strategy Projects of Guangdong Province (2019B020203002), the Basic Research Project of Haixi Agriculture and Animal Husbandry Bureau (Qinghai Caidamu Cashmere Goat Genomic Breeding project), the National Natural Science Foundation of China (31872560 and 31672399), the Shenzhen Municipal Government of China (JCYJ20180307163440037).

We thank Bayaertu & Yong quan from Dongwuzhumuqin Banner Aogali Animal Husbandry Co., Ltd and Hu Yeyong from Henan yudong animal husbandry co. Ltd for providing UC and MT goat materials, respectively. We thank Shancen Zhao for helpful discussion and revising the manuscript.

## Author contributions

L.Y, F.M, Z.S.Y. conceived, designed, and supervised the study. H. H., F. M., L.Y., C.T. and M.Z.P performed the informatics analysis of the sequencing data. Z.S.Y., Y.M.M., W.Q., Y.C.Y., W.Q.J., WL.B., G.G., Mengkedala., D.W.D., L.B., Z.Q.F. obtained goat material and DNA for re-sequencing. Z.S.Y., Y.M.M., W.Q.J., WL.B., G.G., Mengkedala. and L.B. performed phenotypic data collation and analysis. Y.M.M., W.R., J.D., W.Q. and Z.T.T. participated in the laboratory work. Z.X.J., Z.T.T. and L.L. performed dual-Luciferase Reporter analysis. C.T analyzed the transcriptome. L.Y., H.H., Y.M.M., J.D. and Z.X.J are the major contributors in writing the manuscript. All authors read, revised, and approved the final manuscript.

## Additional information

Supplementary Information accompanies this paper at Supplementary.docx and Supplementary table s.xlsx.

