## Supplementary for "Whole-genome sequencing of Chinese native goat offers biological insights into cashmere fiber formation"

### Population sequencing of Chinese native goat reveals an insulator deletion in LHX2 affecting cashmere yield and quality

#### Supplementary Figure List

#### Supplementary Notes 1

##### 1.1 Sample information and sequencing

All 120 goats of 8 breeds were used in this study, which includes 30 Chaidamu Cashmere Goats (CDMC) from Chaidamu City of Qinghai Province; 11 Ujumqin Cashmere goats (UC) from Ujimqin Banner of Inner Mongolia Province; 8 Guizhou Black goats (GZB) from Guiyang City of Guizhou Province; 11 Yunnan Black Bone Goats (YNBB) from Lanping County of Yunnan Province; 10 Matou Goats (MT) from Shangqiu City of Henan Province; 10 Jining Grey Goats(JNG) from Jining City of Shandong Province; 10 Jintang black goat(JTB) from Jintang County of Sichuan Province; 12 Chengdu brown goat(CDB) from Chengdu City of Sichuan Province.

Firstly, ear tissues or blood of goats were collected on-site and stored in alcohol sampling tubes or blood collection tubes; then genomic DNA was extracted by commercial kits, and samples with DNA integrity and concentration > 50ng/ul were used for library construction. Next, the standard circularization step of DNBSEQ was carried out to make DNA nanoballs. Finally, the libraries were sequenced using BGIseq500 with PE100[1].

##### 1.2 Quality control and reads mapping

Quality control of all raw reads was performed using SOAPnuke1.5.0 (https://github.com/BGI-flexlab/SOAPnuke), with the following criteria: quality value ≤20; low-quality bases >30%; and N bases >5%. Clean reads were then mapped to the reference genome ARS1 (https://www.ncbi.nlm.nih.gov/genome/10731?genome_assembly_id=281266) using BWA-0.7.12[2] (https://sourceforge.net/projects/bio-bwa/) with default parameters. In the mapping process, a BAM file index was built using SamTools (https://github.com/samtools/samtools/releases/), followed by the bam file being sorted and removed for duplicated reads with picard-tools-1.105 (https://github.com/broadinstitute/picard/releases).

##### 1.3 Variant calling and annotation

After mapping, the “HaplotypeCaller”, “CombineGVCFs” and “GenotypeGVCFs” in GATK4 were used to detect SNPs and indels with default parameters. The output VCF File was then screened for SNPs using "SelectVariants function" of GATK4. We further reserved the SNPs using "VariantFiltration" of GATK4 using the following criteria: QD < 2.0; FS > 60.0; MQ < 40.0; and ReadPosRankSum < -8.0. Next, we used VCFtools to obtain high-quality SNPs combined with population information. SNPs that met at least one of the following standards were excluded: (i) minor allele frequency (MAF) <0.05; (ii) SNP call rate <90%; and (iii) $P < 0.000001$ for the Hardy–Weinberg equilibrium. Only the loci of two alleles were retained for the subsequent analyses. Finally, ~13 million SNPs were remained for further analysis.

#### Supplementary Notes 2: Population structure and phylogenetic analysis

##### 2.1 Geographic distribution mapping

First, we obtained the longitude and dimension data of 10 goat sampling sites through Google map; then used package "mapdata" and "ggrepel in R for mapping; finally, the Photoshop software was used for organization and optimization.

##### 2.2 Principal component analysis (PCA)

The PLINK was used for principle component analysis (PCA) with 120 goats based on the filtered SNPs using function "--vcf vcf --out pca –pca --chr-set 29 --allow-extra-chr". The contribution of PCA1 and PCA2 was 15.94% and 10.3% respectively. The PCA further reveals that the difference between North China (cashmere goats, JNG and MT) and South China (JTB, CDB, YNBB and GZB) is greater than that between cashmere goats and ordinary goats (**Supplementary Fig. 1**).

##### 2.3 Population admixture

Population structure was inferred using a Bayesian-based approach via the software package Admixture_linux-1.23 (https://dalexander.github.io/admixture/index.html); then, the cross validation was performed to choose the optimal *k* value. The analysis process includes the following three steps:

(1) Converting VCF format to PLINK format. The Linux command line was "vcftools --vcf vcf --plink --out goat.plink";

(2) Using PLINK for further filtering with Linux command line "plink --noweb --file plink --geno 0.05 --maf 0.05 --hwe 0.0001 --chr-set 29 --make-bed --out QC"; after this step we get the corresponding bed file.

(3) Using ADMIXTURE v1.23 for population structure analysis with k values ranged from 3 to 7. Five-fold cross-validation was used to evaluate the fitness. For each k, we ran the ADMIXTURE for 20 times and calculated the corresponding cross-validation errors with the Linux command line "admixture --cv QC.bed $k | tee log${k}.out"; then the results were plotted using R.

**2.4 Phylogenetic tree**

The VCF files were converted into a matrix using VCF2Dis-1.09 (https://github.com/BGI-shenzhen/VCF2Dis), and tree files were formed using PHYLIPNEW-3.69.650 (https://evolution.genetics.washington.edu/phylip.html). Then a phylogenetic tree based on genetic distance was constructed via the neighbor-joining method by iTOL[3] (http://itol.embl.de). The Linux command line of converting to a matrix is "VCF2Dis -InPut vcf -OutPut p_dis.mat"; the Linux command line of constructing a phylogenetic tree is "PHYLIPNEW-3.69.650/bin/fneighbor -datafile p_dis.mat -outfile goat.tree.txt -matrixtype s -treetype n -outtreefile goat.n.tree.tre". The neighbor-joining (NJ) tree revealed strong clustering of the Chinese native goat into three genetic groups that presented a low level of genetic differentiation, which recapitulated the finding in the PCA (Supplementary Fig. 4).

**2.5 LD decay**

PopLDdecay (https://github.com/BGI-shenzhen/PopLDdecay) was used to calculate the $r^2$ of linkage disequilibrium (LD) with command "pLDdecay -InVCF vcf -MaxDist 300 -OutStat LDdecay".

#### Supplementary Notes 3: Gene flows and effective population size

##### 3.1 Gene flow

A population-level admixture analysis was carried out with TreeMix (version 1.12) program (Pickrell and Pritchard 2012). It uses genome-wide SNP sites to infer the Maximum Likelihood (ML) tree for the 12 goat breeds (236 individuals) and an outgroup (Bezoars), with the command '-i input -bootstrap -k 500 -root IRW -o output'. The covariance matrix of the ML tree was estimated to reflect the correspondence between the ML tree and the SNP data and to identify pairs of populations that showed poor fits in the ML tree. When the ML tree did not fully describe the data, an admixture analysis was performed by allowing migration events. The poor-fit populations were regarded as candidates for adding potential migration edges. Migration events are gradually added to the ML trees until the model can explain 98% of the variation in breeds. The command was '-i input -bootstrap -k 500 -m migration events -o output'. The maximum-likelihood (ML) tree without migration events inferred from the TreeMix analysis divided the 236 goats into three clusters, which is similar to the population structuring patterns identified from the phylogenetic tree, PCA and genetic structure analysis (**Supplementary Fig. 5**). Migration events were detected among the three clusters when potential migration edges were added to the ML tree (**Supplementary Fig. 5**).

##### 3.2 Effective population size

PSMC software was used to infer historical dynamics of effective population size and the divergence timing of goats based on seven samples from four lineages with a high sequencing depth[4]. The whole-genome diploid consensus sequences for each sample were generated by SAMtools and BCFtools with the parameter C50. Sites with sequencing depths < 10 and > 100 (vcfutils.pl vcf2fq -d 10 -D 100) were removed to reduce the probability of false positives. The PSMC parameter (psmc -N25 -t15 -r5 -p "4+25*2+4+6") was used to infer the historical effective population size, where the parameters of generation time and mutation rate were set to 2.0 and $2.5e^{-8}$, respectively.

#### Supplementary Notes 4: Selective sweep analysis

Fst and θπ were performed to evaluate fixation and differentiation. High-quality SNPs were chosen to calculate Fst using VCFtools based on sliding 150kb windows with a 10kb step length. Fst values were then standardized as ZFst by calculating Z-scores according to the formula $x^*=(\bar{x}-x)/\sigma$. We also calculated the $\theta_\pi$ ratios and $\log_2$-transformed, namely $Log_2(\theta_\pi$ Ratios). We annotated genes functions for those windows with larger ZFst and $Log_2(\theta_\pi$ Ratios) values (top 1%) [5,6]. Gene ontology and KEGG analyses were performed using Metascape[7] and compared to the human genome as background.

##### 4.1 Genome-Wide Selective Sweep between cashmere and ordinary goat

We performed selective sweep analysis with genome sequences of 84 cashmere goats (IMC, CDMC, LNC) and 58 ordinary goats (YNBB, GZB, JTB and GZB) We choose those windows with top 1% of statistic (ZFst>4.042 and $\log_2(\theta_\pi$ Ratios)> 1.348) for candidate windows. Finally, we identified 141 candidate genes in and around these windows (500kb from both sides) (Supplementary Table s2 and Supplementary Fig. 6).

##### 4.2 Genome-Wide Selective Sweep for hair color

First, we scanned for selection signatures for the white coat group (CDMC, UC, IMC, LNC) and the black coat group (YNBB, GZB). We identified signature regions harboring *KIT*, *KITG* and *IRF4*(Supplementary Fig.7 and Supplementary Table s4) and 100 kb copy number variants (CNVs) downstream the *KIT* (chr6: 70,859,258-70,959,918) (Supplementary Fig. 10) , which is consistent with the previously report [8]. We next searched for the selection signature in the white coat group (LNC, IMC, CDMC and UC) and the brown coat group (CDM),We also identified a striking signature harboring *ASIP*, *AHCY* and *ITCH* (Supplementary Fig. 8-9 and Supplementary Table s3) and ~154 kb CNV partly overlapping with *ASIP* (chr13:63,334,008-63,380,258) (Supplementary Fig. 11), which is consistent with the previously reported [8].

#### Supplementary Notes 5: Deletion of breakpoints of *LHX2* and *FGF5*

To detect the heterozygous genotype of goats more accurately, we designed PCR primers to detect the deletions in *LHX2* and *FGF5* (Supplementary Table 8 and Supplementary Table 9). Also, we used the same primers to detect the deletions in several sheep breeds (Hu sheep, Ujumqin sheep, and Australian white sheep) (Supplementary Fig. 21).

#### Supplementary Figures

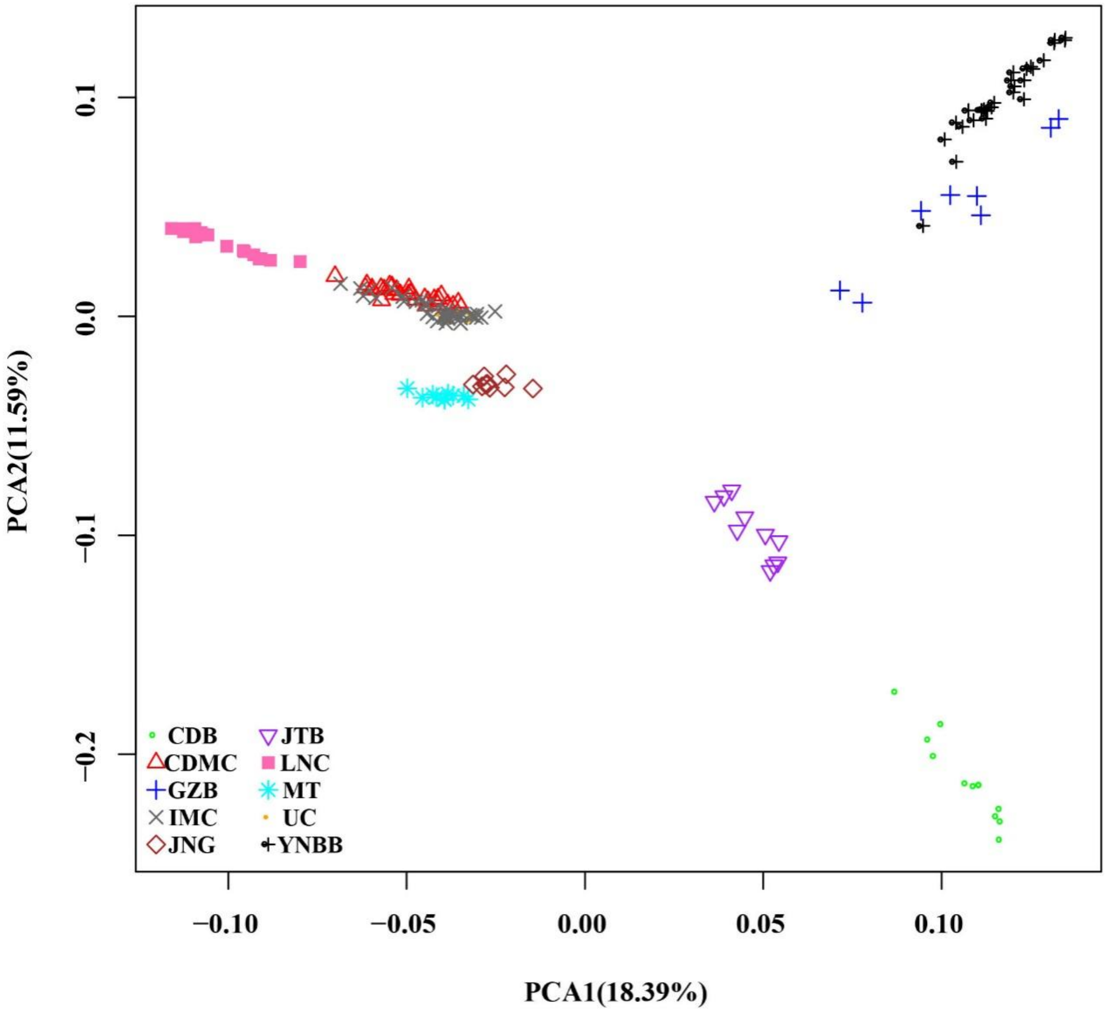

Supplementary Fig. 1 PCA analysis.

Principal components 1 and 2 of 10 local native goat breeds. The horizontal axis represents principal component 1 (PCA1), and the vertical axis represents principal component 2 (PCA2). Four cashmere goat breeds (CDMC, IMC, UC, LNC) were gathered; JNG and MTG breeds are close to the cashmere goats; The JTB, CDB, YNBB and GZB in Southwest China were all far from the cashmere goats and were divided into two small groups.

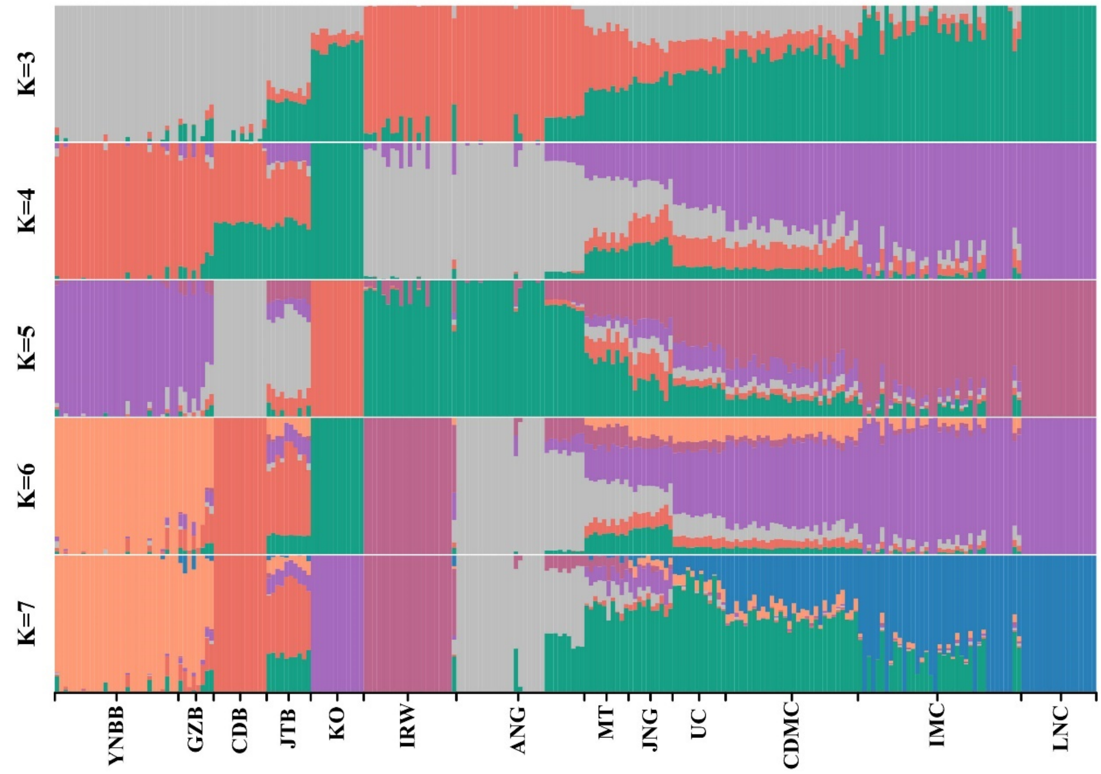

Supplementary Fig. 2 Results of genetic structure analysis of 13 goat breeds.
The length of each colored segment represents the proportion of the individual's genome inferred from K=3-7 ancestral populations. For the abbreviations of the individuals see supplementary table 3-5.

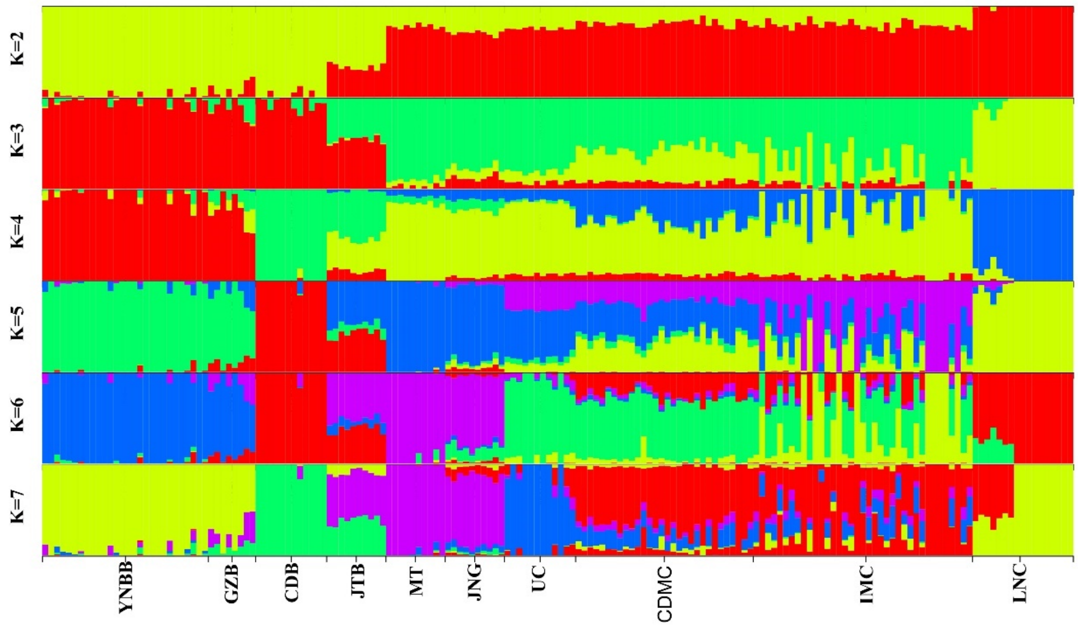

Supplementary Fig. 3 Results of genetic structure analysis of 10 Chinese goat breeds. The length of each colored segment represents the proportion of the individual's genome inferred from K=2-7 ancestral populations. For the abbreviations of the individuals see supplementary table 3-4.

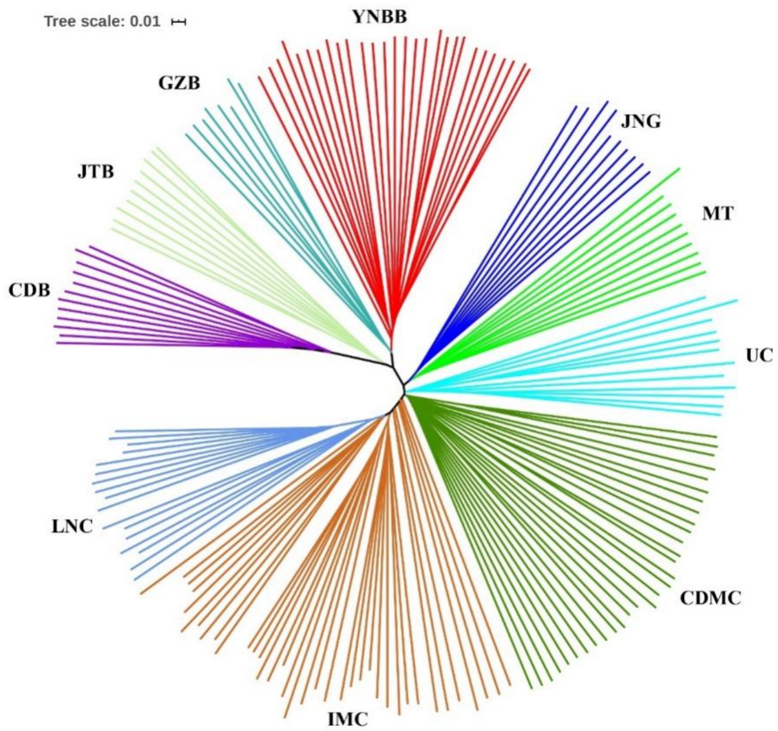

Supplementary Fig. 4 Phylogenetic tree of 10 Chinese native goat breeds.

Each color represents a breed, and each line represents an individual. Ten local goat breeds were clustered into two groups: four goat breeds in southwest China (CDB, JTB, GZB and YNBB) clustered for one group; LNC, IMC, CDMC, UC, MT and JNG gathered for another, JNG and MTG have clustered separately close to the cashmere goat subgroup.

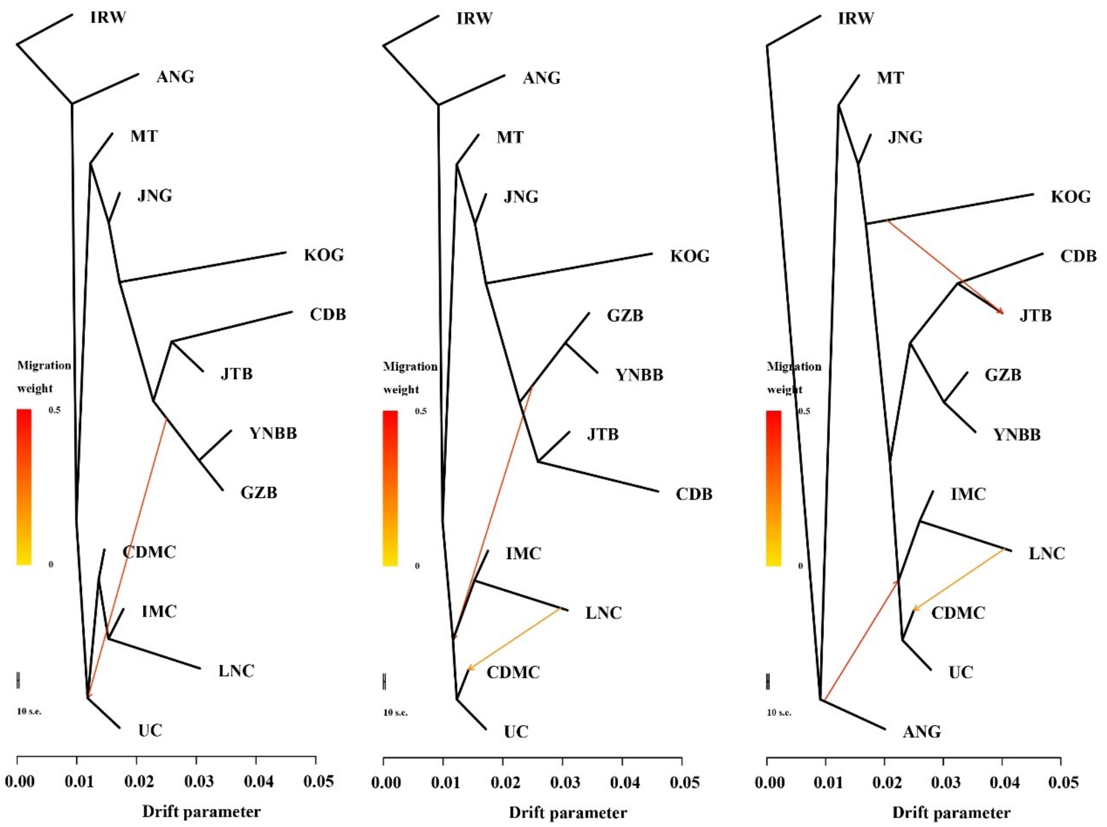

Supplementary Fig. 5 Inferred goat tree of mixture events deduced by TreeMix.
IRW samples were assigned as "outgroup" to root the tree. The graph inferred for the goat populations, allowing one, two and three migration events. Migration arrows are colored according to their weight. Horizontal branch lengths are proportional to the amount of genetic drift that has occurred on each branch. The scale bar shows ten times the average standard error of the entries in the sample covariance matrix.

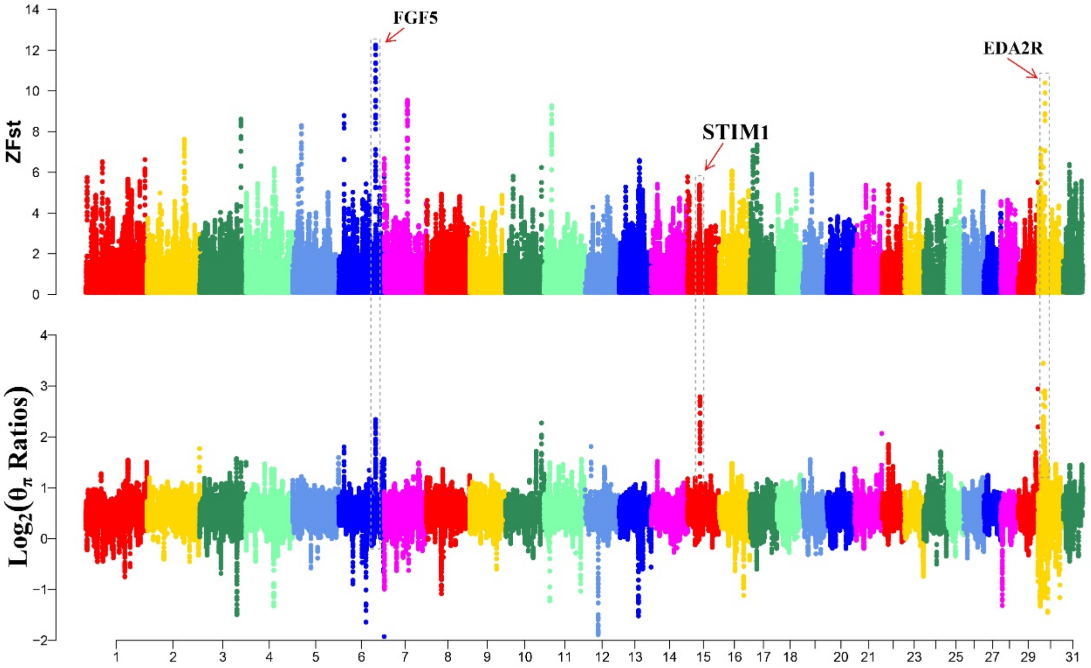

Supplementary Fig. 6 Analysis of the signatures of positive selection in the genome of cashmere and ordinary goat breeds.

Manhattan plot of the genome-wide distribution of pairwise ZFst and $\log_2(\theta_\pi$ Ratios) between cashmere goats (CDMC, IMC, LNC) and ordinary goats (YNBB, GZB, JTB, CDB) using a 150 kb window size and a 10 kb step size. The key candidate genes *FGF5, STIM1, KIT* related to cashmere were annotated at the significant signal. Red arrows and names indicate the top three candidate genes.

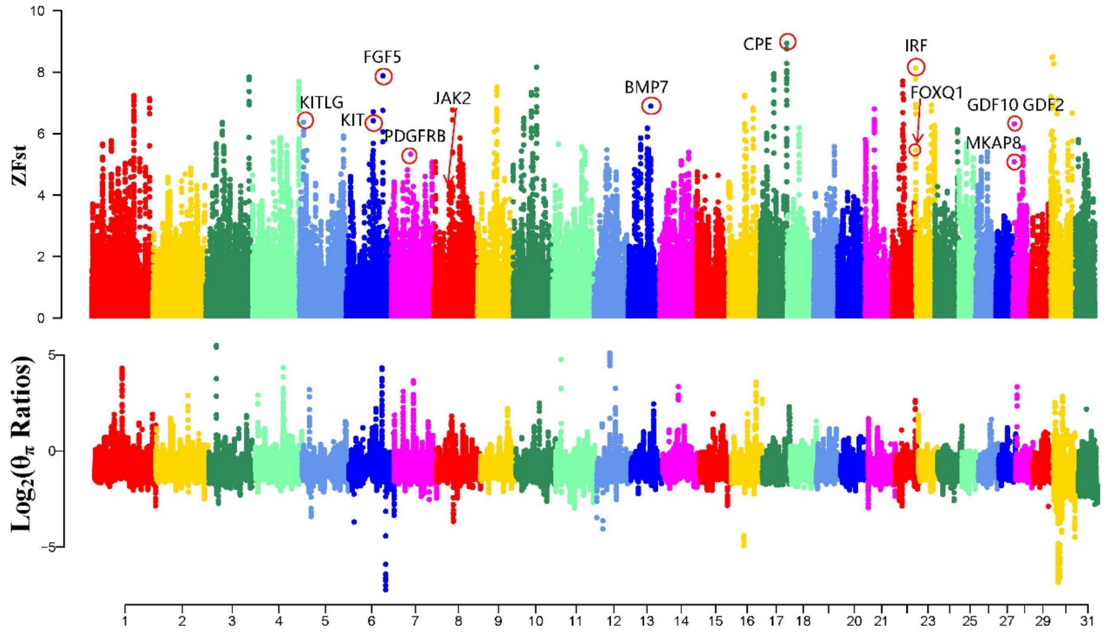

Supplementary Fig. 7 Analysis of the signatures of positive selection in the genome of goat breeds with white and black coat color.

Manhattan plot of the genome-wide distribution of pairwise ZFst and $\log_2(\theta_\pi$ Ratios) between white coat color goats (UC, CDMC, IMC, LNC) and black coat color goats (YNBB, GZB) using a 150 kb window size and a 10 kb step size. The key candidate genes *KIT*, *KITLG*, *FGF5*, *JAK2*, *FOXQ1* related to cashmere were annotated at the significant signal. Red circles and names indicate the top three candidate genes.

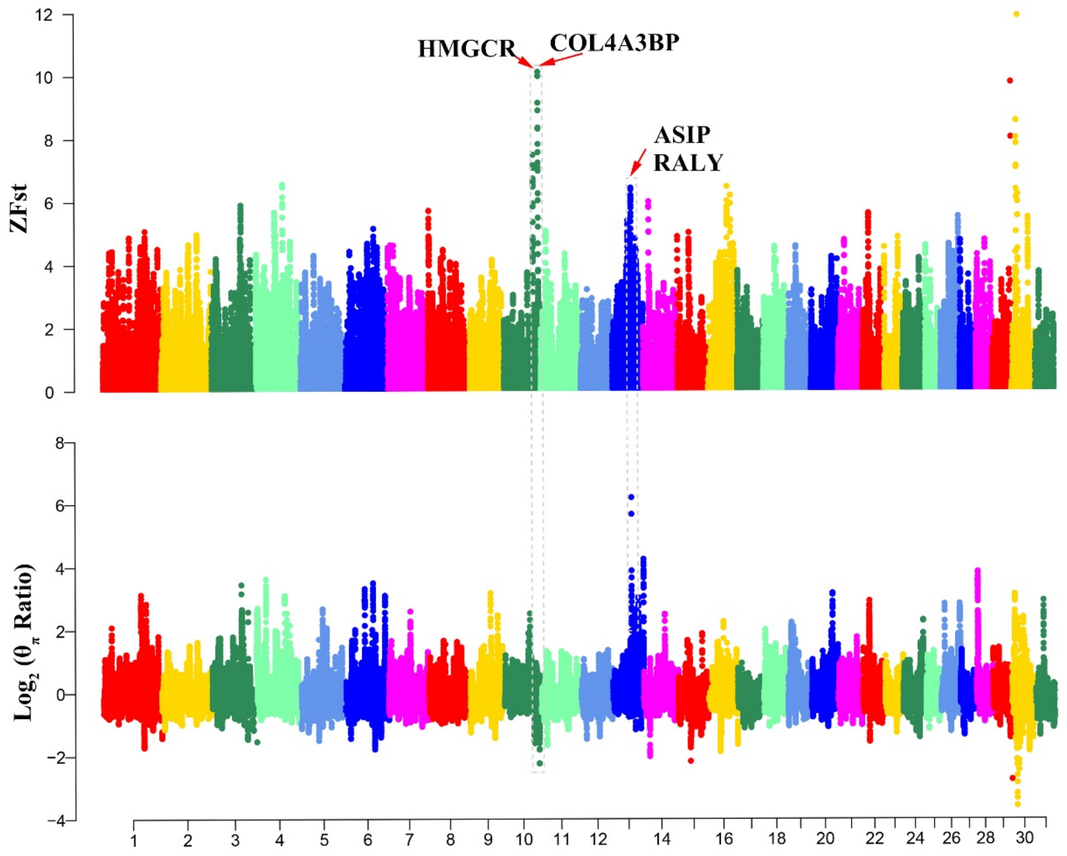

Supplementary Fig. 8 Analysis of the signatures of positive selection in the genome of goat breeds with white and brown coat color.

Manhattan plot of the genome-wide distribution of pairwise ZFst and $\log_2(\theta_\pi$ Ratios) between white coat color goats (UC, CDMC, IMC, LNC) and brown coat color goats (CDB) using a 150 kb window size and a 10 kb step size. The key candidate genes *ASIP*, *RALY*, *HMGCR*, *COL4A3BP* related to cashmere color were annotated at the significant signal. Red arrows and names indicate the top three candidate genes.

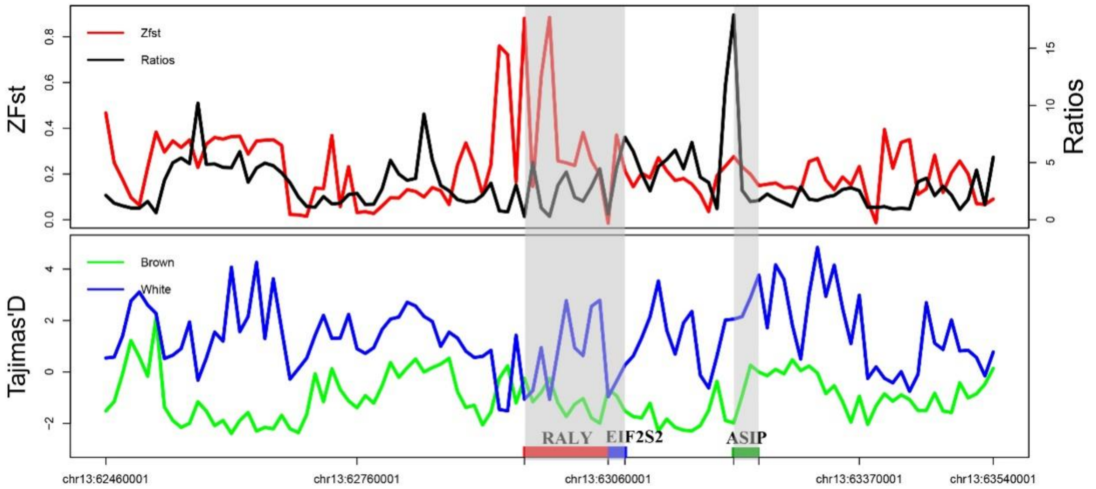

Supplementary Fig. 9 Ratios and ZFst values around the *ASIP* locus.

The black and red lines represent θπ Ratios and ZFst values, respectively. Tajima's D values around the *RALY*, *EIF2S2*, *ASIP* locus. The green and blue lines represent the brown coat color goats (CDB) and the white coat color goats (CDMC, IMC, UC, LNC).

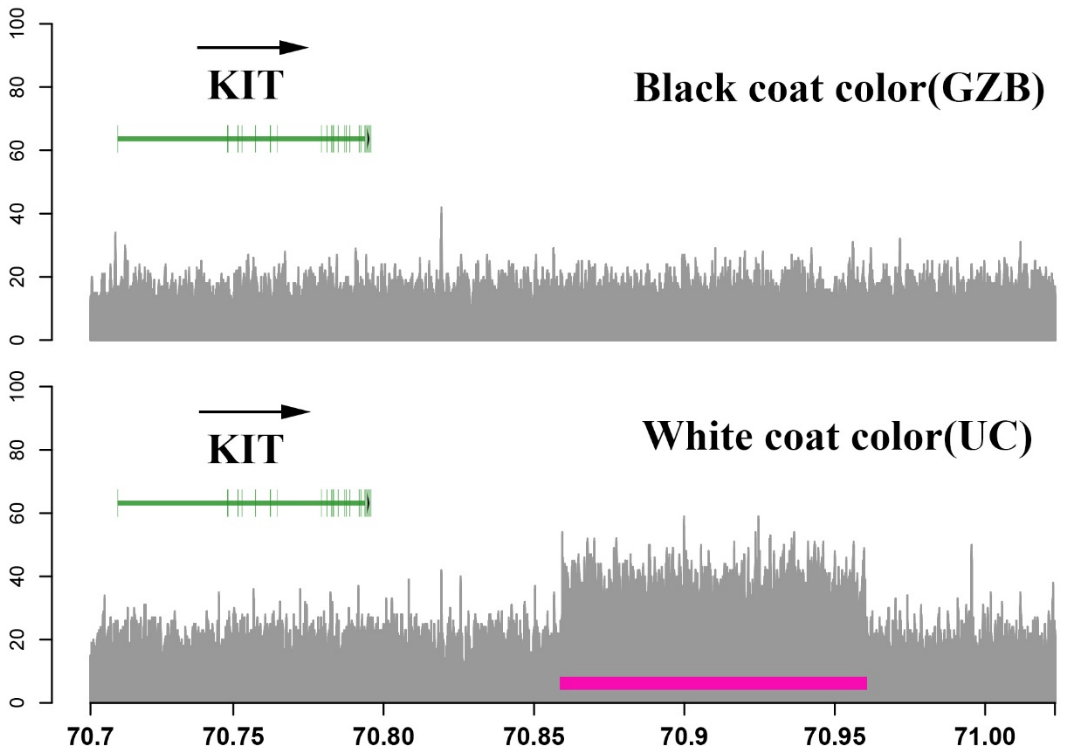

Supplementary Fig. 10 CNVs at the *KIT* locus.

The black coat color goat (GZB) does not show copy number variation (upper panel); however, in the white coat color goat (UC), the coverage plot shows a ~100 kb duplicated CNV (chr6:70859258-70959918) downstream of the *KIT* gene (lower panel).

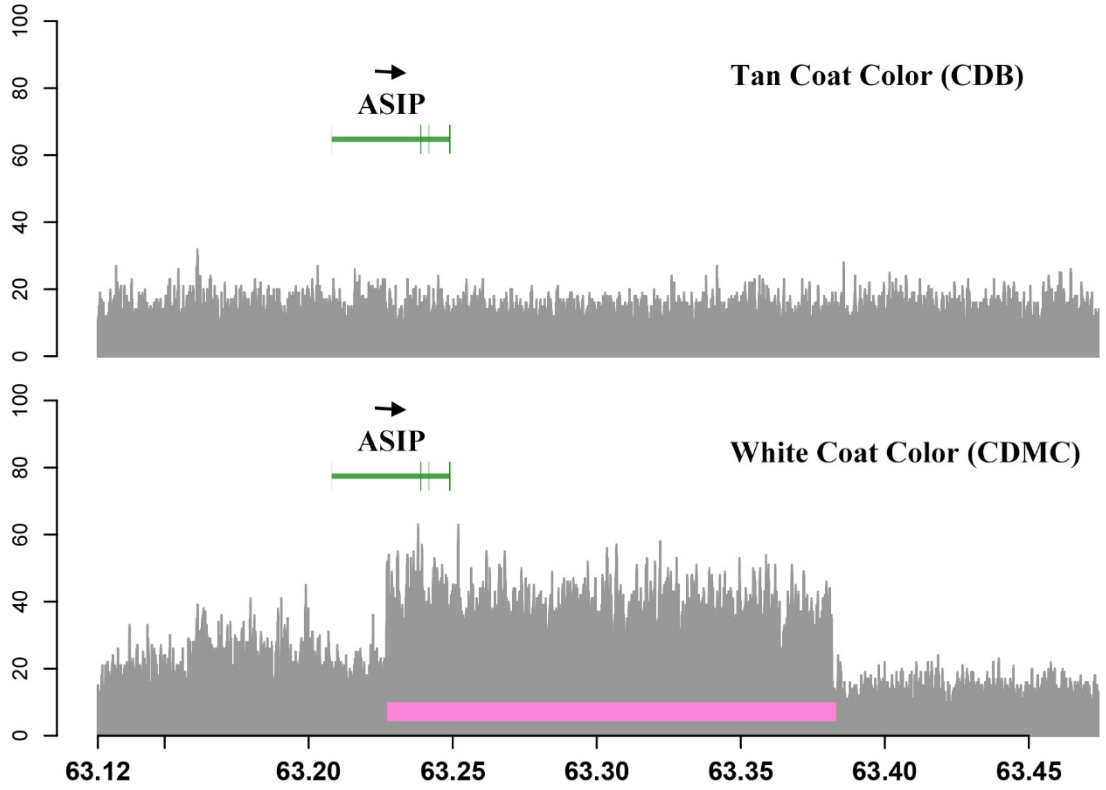

Supplementary Fig. 11 CNVs at the ASIP locus.
The black coat color goat (CDB) does not show any copy number variation. However, in the tan coat color goat (CDMC), the coverage plot shows a duplication of ~154 kb(chr13:63226824-63381501) downstream of the *KIT* gene.

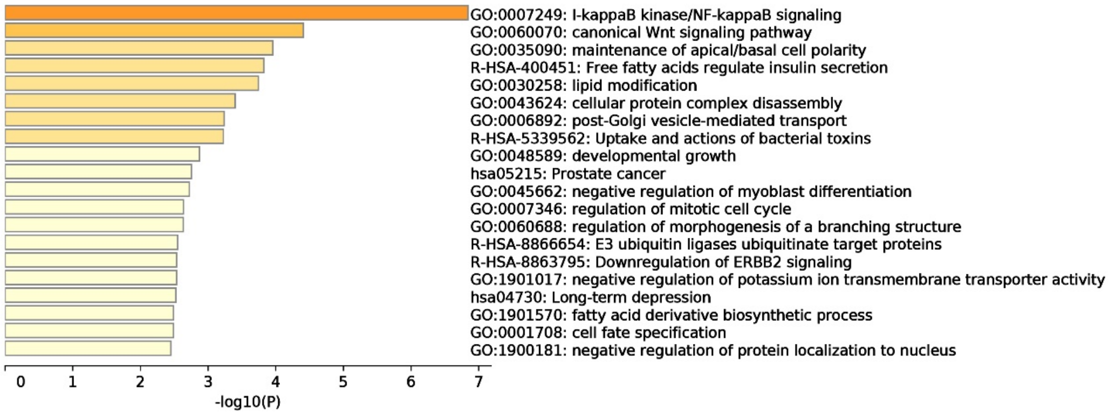

Supplementary Fig. 12 Bar graph of enriched terms across positively selected genes with GO term enrichment analysis.
X-axis represents the p-values.

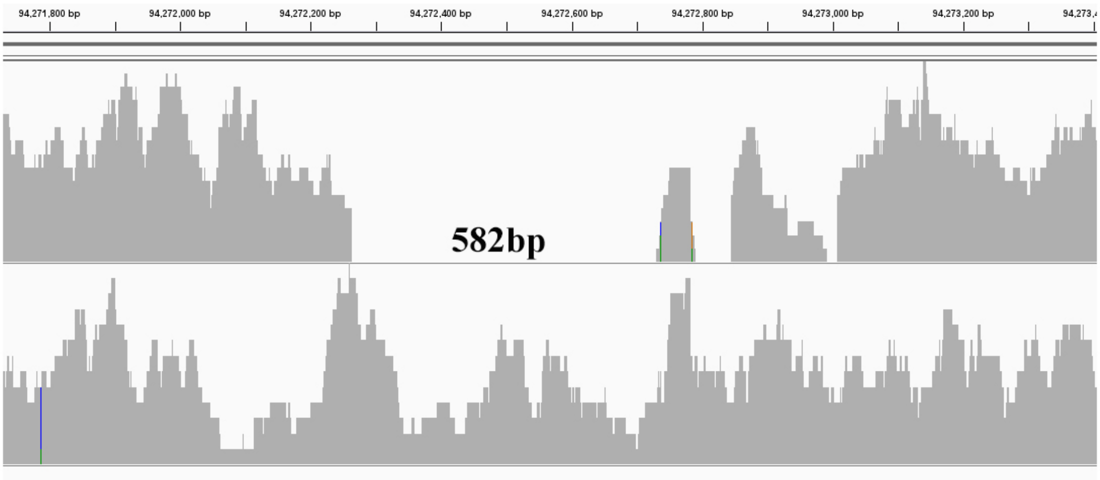

Supplementary Fig. 13 Visualization of whole genome sequencing data identifies 582 bp deletion in the ancient goat.

BAM files were visually inspected using the Integrative Genomics Viewer (IGV).

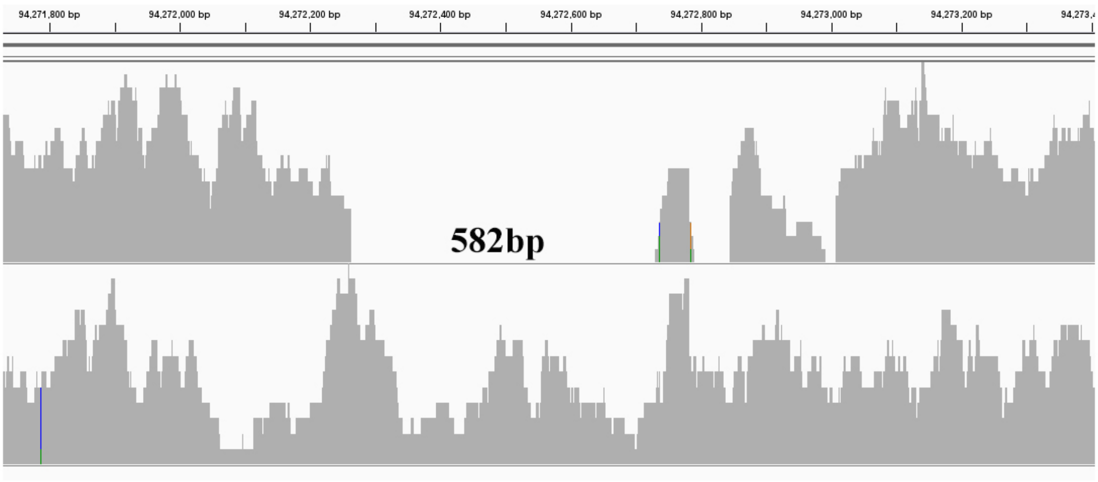

Supplementary Fig. 14 Visualization of Whole Genome Sequencing Data identifies 582 bp deletion near *LHX2* in the ibex goats.

Upper panel represents ibex goat carries 582bp deletion and lower panel not carries.

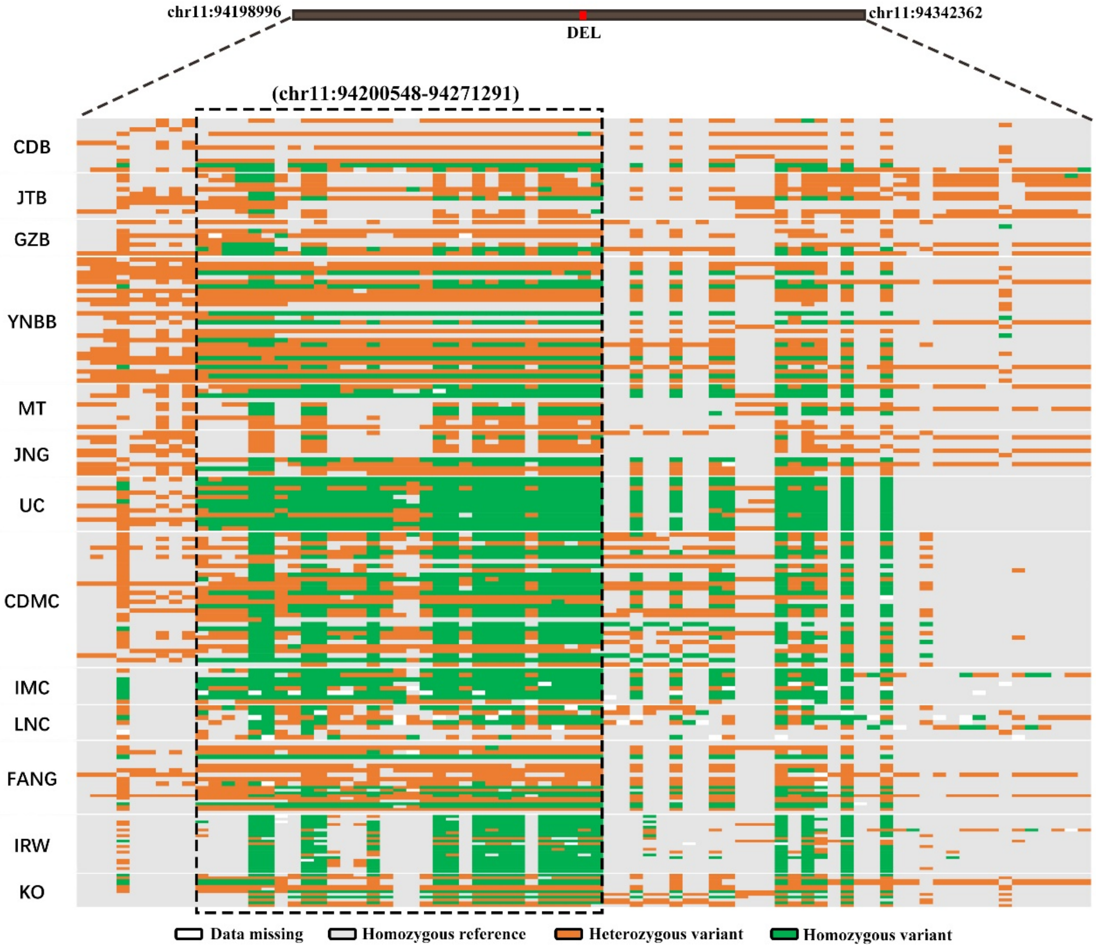

Supplementary Fig. 15 The pattern of SNP genotypes near DEL among 13 goat breeds.

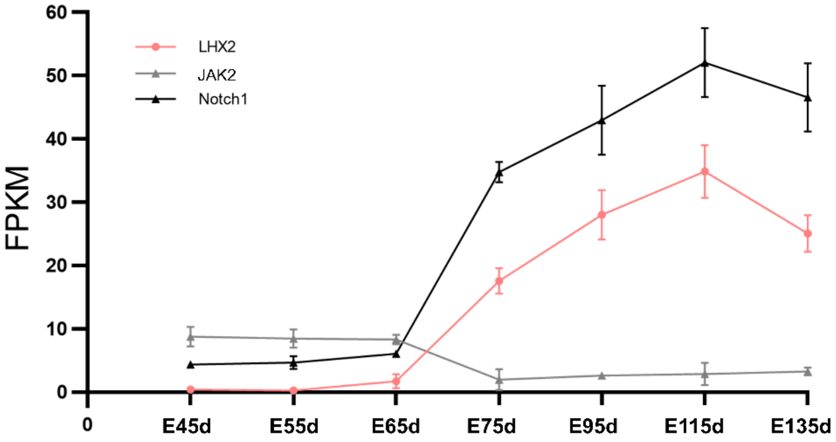

Supplementary Fig. 16 The expression of *LHX2*, *JAK2* and *Notch1* in different days (from 45 d to 135 d) of fetal skin.

E represents days of gestation. The expression data is downloaded from the published data[9].

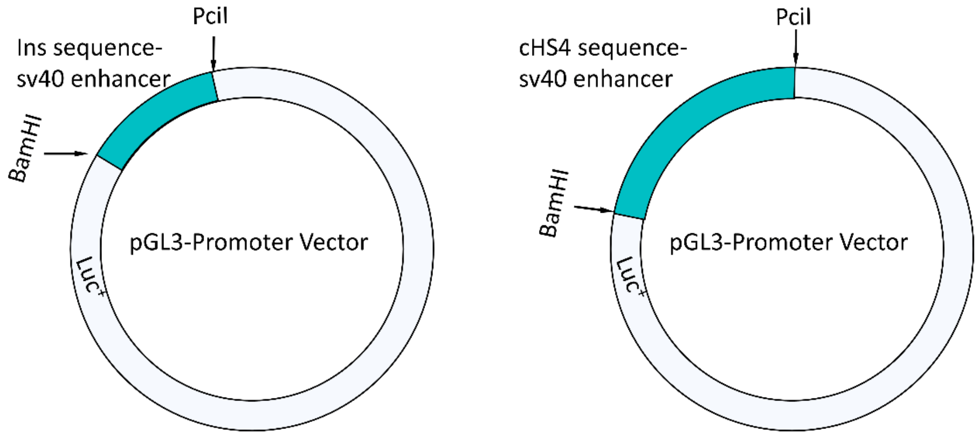

Supplementary Fig. 17 Schematic drawing of the construction of DNA plasmids.
The 796 bp DNA fragment (including 582 bp deletion sequence and some flanking sequence, named Ins and cHS4) and subsequently inserted downstream and upstream of the SV40 promoter in pGL3 plasmid.

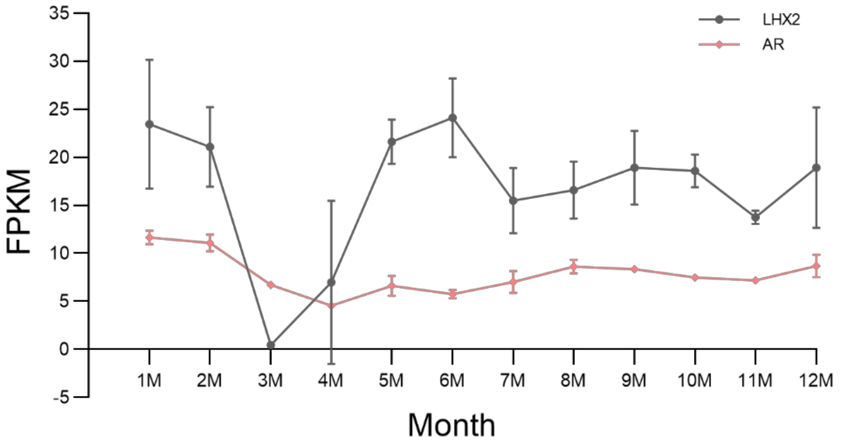

Supplementary Fig. 18 The expression of *LHX2* and *AR* in different months of a year. The expression data is downloaded from the NCBI SRA database[10] (PRJNA470971).

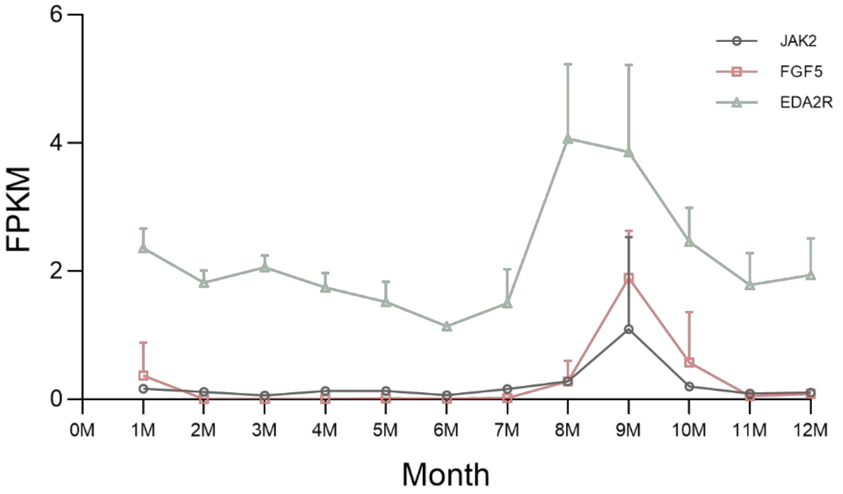

Supplementary Fig. 19 The expression of *JAK2* and *EDA2R* as well as *FGF5* in different months of a year.

The expression data is downloaded from the NCBI SRA database[10] (PRJNA470971).

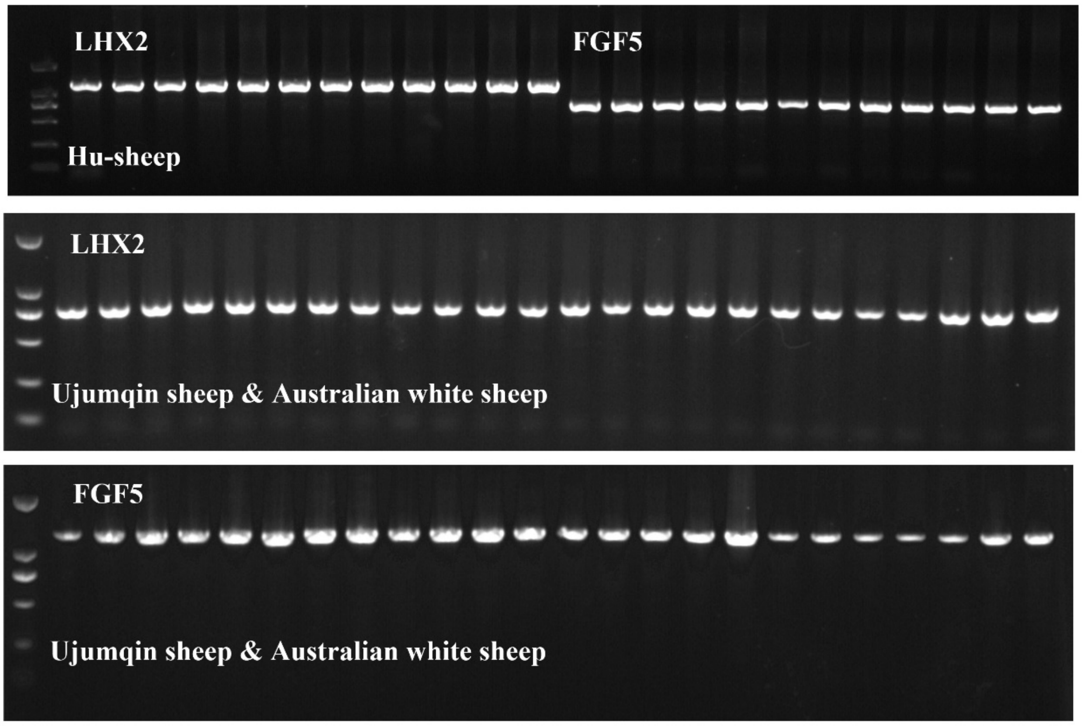

Supplementary Fig. 20 The PCR identification of deletions in sheep.

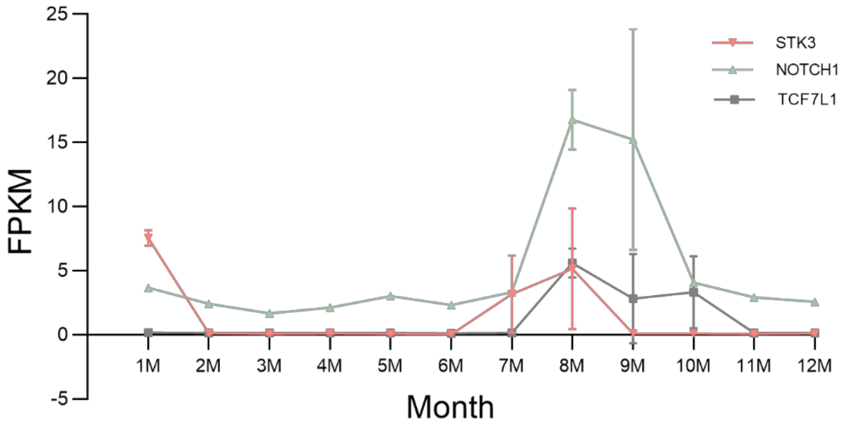

Supplementary Fig. 21 The expression of *STK2* and *NOTCH1* as well as *TCF7L1* in different months of a year.

The expression data is downloaded from the NCBI SRA database[10] (PRJNA470971).

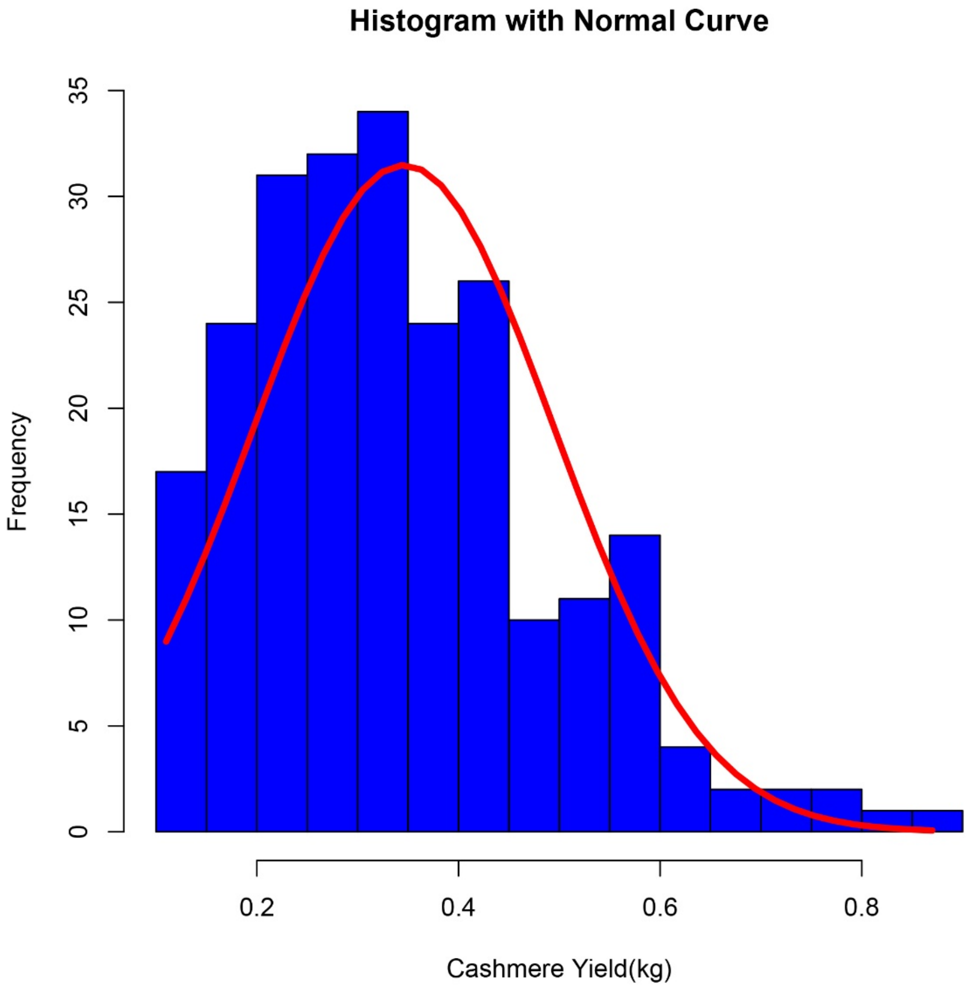

Supplementary Fig. 22 Distribution diagram of cashmere yield data

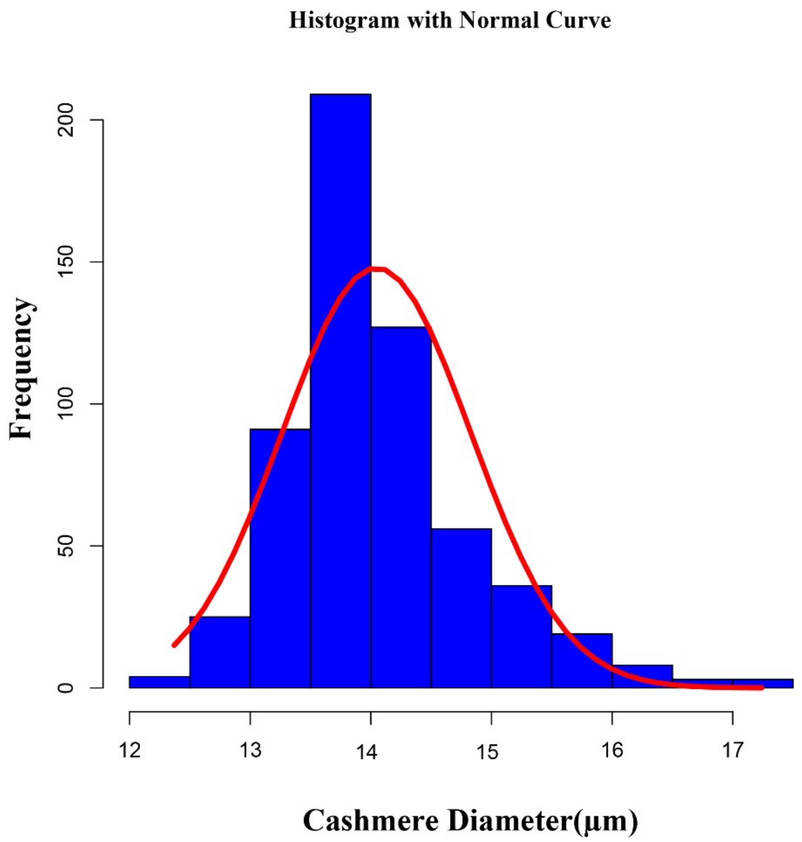

Supplementary Fig. 23 Distribution of cashmere diameter measurements

### Supplementary Tables

Supplementary Table 1 Overview of sequencing information of 120 goats

| Abbre viation | Sampling location | ID | Size_MB | Depth | Coverage | Mapping_rate(%) |
|---|---|---|---|---|---|---|
| CDMC | Chaidamu, Qinghai | HXR0157 | 24722.84 | 8.09 | 99.65% | 99.19 |
| | | HXR0014 | 27535.99 | 9.33 | 99.70% | 99.21 |
| | | HXR0138 | 28576.42 | 9.43 | 99.72% | 99.2 |
| | | HXR0044 | 29417.06 | 10.04 | 99.71% | 99.22 |
| | | HXR1167 | 29859.92 | 10.18 | 99.68% | 99.36 |
| | | HXR0147 | 31318.85 | 10.51 | 99.72% | 99.19 |
| | | HXR0047 | 31448.91 | 10.46 | 99.73% | 99.2 |
| | | HXR1188 | 32735.31 | 11.07 | 99.71% | 99.28 |
| | | HXR1166 | 32867.20 | 11.27 | 99.71% | 99.3 |
| | | HXR171 | 34456.79 | 11.82 | 99.72% | 99.26 |
| | | HXR0981 | 34904.03 | 11.86 | 99.69% | 99.24 |
| | | HXR1321 | 30607.06 | 10.28 | 99.54% | 99.17 |
| | | HXR1750 | 26659.97 | 9.09 | 99.55% | 99.22 |
| | | HXR2752 | 28146.61 | 9.53 | 99.60% | 99.22 |
| | | HXR0908 | 28744.20 | 9.94 | 99.66% | 99.27 |
| | | HXR1630 | 33845.47 | 11.68 | 99.72% | 99.09 |
| | | HXR3386 | 30860.89 | 10.67 | 99.65% | 99.25 |
| | | HXR3388 | 30735.56 | 10.6 | 99.65% | 99.27 |
| | | HXR0022 | 26253.44 | 8.7 | 99.58% | 99.24 |
| | | HXR0045 | 29850.22 | 10.09 | 99.63% | 99.3 |
| | | HXR0126 | 28236.77 | 9.4 | 99.67% | 99.27 |
| | | HXR0170 | 28779.79 | 9.6 | 99.65% | 99.28 |
| | | HXR0186 | 27155.72 | 9.06 | 99.64% | 99.25 |
| | | HXR0370 | 33842.07 | 11.2 | 99.73% | 99.18 |
| | | HXR0711 | 34055.78 | 11.64 | 99.71% | 99.27 |
| | | HXR097 | 30425.80 | 10.06 | 99.68% | 99.26 |
| | | HXR0127 | 36488.06 | 12.6 | 99.72% | 99.27 |
| | | HXR0176 | 29880.23 | 10.19 | 99.65% | 99.28 |
| | | HXR2781 | 35434.27 | 12.17 | 99.71% | 99.28 |
| | | HXR3030 | 31065.93 | 10.6 | 99.68% | 99.17 |
| UC | Ujumqin,Inner Mongolia | WMG0010 | 29808.94 | 10.22 | 99.68% | 99.22 |
| | | WMG0013 | 25158.27 | 8.53 | 99.58% | 99.19 |
| | | WMG0016 | 32235.68 | 10.99 | 99.61% | 99.33 |
| | | WMG0017 | 32123.61 | 11.03 | 99.70% | 99.23 |
| | | WMG0004 | 45279.43 | 15.93 | 99.76% | 99.23 |

| | | | | | | |
|---|---|---|---|---|---|---|
| | | WMG0005 | 37758.79 | 13 | 99.73% | 99.25 |
| | | WMG0006 | 43441.20 | 15.16 | 99.73% | 99.22 |
| | | WMG0007 | 35369.19 | 12.18 | 99.73% | 99.23 |
| | | WMG0008 | 39552.82 | 13.77 | 99.76% | 99.24 |
| | | WMG0009 | 30251.41 | 10.41 | 99.71% | 99.21 |
| | | WMG0011 | 29613.21 | 10.01 | 99.70% | 99.23 |
| | | WMG0012 | 39691.25 | 13.55 | 99.74% | 99.24 |
| CDB | Chengdu,Sichuan | CMS0019 | 28356.30 | 9.82 | 99.66% | 99.25 |
| | | CMS0084 | 34839.14 | 12.21 | 99.69% | 99.27 |
| | | CMS0177 | 34653.57 | 12.04 | 99.69% | 99.4 |
| | | CMS0330 | 32826.68 | 11.55 | 99.67% | 99.38 |
| | | CMS0037 | 31909.39 | 10.83 | 99.69% | 99.28 |
| | | CMS0088 | 39151.30 | 13.21 | 99.70% | 99.26 |
| | | CMS0100 | 28772.64 | 9.75 | 99.61% | 99.39 |
| | | CMS0406 | 27437.51 | 9.34 | 99.63% | 99.28 |
| | | CMS0026 | 33172.37 | 10.83 | 99.66% | 99.3 |
| | | CMS0053 | 31938.17 | 10.64 | 99.67% | 99.32 |
| | | CMS0065 | 25939.66 | 8.68 | 99.58% | 99.28 |
| | | CMS0431 | 28635.63 | 9.36 | 99.60% | 99.33 |
| GZB | Guiyang,Guizhou | GHS0004 | 16639.80 | 5.39 | 98.76% | 98.98 |
| | | GHS0013 | 20755.17 | 6.93 | 99.46% | 98.99 |
| | | GHS0010 | 24577.96 | 8.2 | 99.60% | 98.56 |
| | | GHS0001 | 26279.23 | 8.89 | 99.65% | 98.88 |
| | | GHS0027 | 27464.94 | 9.22 | 99.68% | 99.18 |
| | | GHS0017 | 27865.07 | 9.4 | 99.64% | 99.1 |
| | | GHS0026 | 29457.34 | 9.86 | 99.70% | 99.22 |
| | | GHS0025 | 37752.37 | 13.01 | 99.75% | 99.18 |
| JTB | Jintang,Sichuan | JTY0038 | 22812.64 | 7.84 | 99.56% | 99.2 |
| | | JTY0076 | 25960.19 | 8.93 | 99.66% | 99.23 |
| | | JTY0042 | 26355.43 | 9.26 | 99.65% | 99.25 |
| | | JTY0001 | 27932.10 | 9.67 | 99.62% | 99.24 |
| | | JTY0045 | 29104.36 | 9.92 | 99.68% | 99.21 |
| | | JTY0022 | 30724.89 | 10.59 | 99.70% | 99.03 |
| | | JTY0062 | 30799.82 | 10.83 | 99.71% | 99.21 |
| | | JTY0012 | 32126.81 | 10.9 | 99.71% | 99.1 |
| | | JTY0034 | 32642.38 | 11.22 | 99.72% | 99.09 |
| | | JTY0050 | 35249.33 | 11.95 | 99.75% | 99.22 |
| YNBB | Lanping,Yunnan | BBG97 | 68903.47 | 24.77 | 99.68% | 99.49 |
| | | BBG274 | 53166.99 | 19 | 99.64% | 99.54 |
| | | BBG3101 | 67090.35 | 23.88 | 99.72% | 99.62 |
| | | BBG6106 | 57484.93 | 20.78 | 99.69% | 99.56 |
| | | BBG7111 | 57943.70 | 20.91 | 99.67% | 99.53 |
| | | BBG8111 | 57948.10 | 20.75 | 99.70% | 99.51 |

| | | | | | | |
|---|---|---|---|---|---|---|
| | | BBG974 | 56834.47 | 20.32 | 99.69% | 99.47 |
| | | BBG10101 | 52573.69 | 18.74 | 99.69% | 99.52 |
| | | BBG12142 | 67912.64 | 24.3 | 99.68% | 99.61 |
| | | BBG1674 | 55152.12 | 19.84 | 99.65% | 99.55 |
| | | BBG1790 | 61289.37 | 21.93 | 99.69% | 99.62 |
| | | BBG1897 | 69885.29 | 24.88 | 99.72% | 99.54 |
| | | BBG26101 | 55940.49 | 20.26 | 99.70% | 99.46 |
| | | BBG28106 | 67676.59 | 24.27 | 99.68% | 99.35 |
| | | BBG31107 | 74517.03 | 26.94 | 99.68% | 99.54 |
| | | BBG3674 | 58077.22 | 20.82 | 99.66% | 99.61 |
| | | BBG39133 | 67997.18 | 24.34 | 99.71% | 99.64 |
| | | BBG8094 | 59123.55 | 21.11 | 99.68% | 99.52 |
| | | BBG8166 | 59744.89 | 21.79 | 99.70% | 99.6 |
| | | BBG8266 | 56908.26 | 20.7 | 99.70% | 99.55 |
| | | BBG8396 | 67991.46 | 24.94 | 99.71% | 99.53 |
| | | BBG8990 | 59468.12 | 21.62 | 99.70% | 99.57 |
| | | BBG90100 | 64959.42 | 23.48 | 99.69% | 99.66 |
| | | BBG9195 | 59900.10 | 21.55 | 99.68% | 99.54 |
| | | BBG93108 | 64680.66 | 11.98 | 99.54% | 99.67 |
| | | BBG9590 | 55525.50 | 20.04 | 99.70% | 99.51 |
| | | BBG9896 | 60156.38 | 21.58 | 99.70% | 99.63 |
| | | BBG99111 | 61696.89 | 22.19 | 99.69% | 99.51 |
| JNG | Jining,Shandong | GRG43 | 80147.53 | 27.97 | 99.69% | 98.74 |
| | | GRG45 | 77785.36 | 26.92 | 99.57% | 98.73 |
| | | GRG47 | 70586.37 | 24.5 | 99.55% | 98.74 |
| | | GRG77 | 67499.73 | 23.42 | 99.64% | 98.76 |
| | | GRG76 | 58886.59 | 20.43 | 99.10% | 98.65 |
| | | GRG75 | 62095.07 | 21.69 | 99.58% | 98.79 |
| | | GRG15 | 74124.89 | 26 | 99.70% | 98.62 |
| | | GRG16 | 64099.62 | 23.07 | 99.68% | 98.74 |
| | | GRG17 | 58185.73 | 20.13 | 99.66% | 98.66 |
| | | GRG18 | 66355.72 | 22.34 | 99.65% | 98.53 |
| MT | Shangqiu,Henan | MTG0003 | 33073.79 | 12.05 | 99.71% | 99.3 |
| | | MTG0005 | 31190.80 | 11.44 | 99.71% | 99.29 |
| | | MTG0006 | 32001.89 | 11.67 | 99.72% | 99.29 |
| | | MTG0007 | 36583.23 | 13.23 | 99.72% | 99.3 |
| | | MTG0012 | 27846.19 | 10.16 | 99.62% | 99.3 |
| | | MTG0013 | 28180.39 | 10.31 | 99.67% | 99.26 |
| | | MTG0001 | 28026.49 | 10.39 | 99.66% | 99.34 |
| | | MTG0002 | 23833.58 | 8.79 | 99.58% | 99.28 |
| | | MTG0009 | 28787.71 | 10.5 | 99.67% | 99.3 |
| | | MTG0010 | 27156.25 | 9.93 | 99.62% | 99.29 |

Supplementary Table 2 Sequencing information of 95 published goats used in this study

| Abbreviation | Sampling location | SRR | ID | Size_Mb | Depth(X) |
|---|---|---|---|---|---|
| IMC | Inner Mongolia | SRR4052649<br>SRR4096727 | IMCG02 | 13352.12 | 5.15 |
| | | SRR4052677<br>SRR4096728 | IMCG05 | 10378.24 | 5.92 |
| | | SRR4053244<br>SRR4096730 | IMCG25 | 20533.36 | 7.12 |
| | | SRR4053410<br>SRR4096705 | IMCG28 | 16367.42 | 6.06 |
| | | SRR4053460<br>SRR4096706 | IMCG29 | 19661.99 | 6.95 |
| | | SRR4053519<br>SRR4096707 | IMCG30 | 17400.57 | 6.37 |
| | | SRR4053642<br>SRR4096720 | IMCG43 | 10480.34 | 4.17 |
| | | SRR4051874<br>SRR4101814 | IMCG21 | 18138.44 | 6.33 |
| | | SRR4101803<br>SRR4051950 | IMCG37 | 12121.82 | 4.73 |
| | | SRR4101805<br>SRR4051959 | IMCG39 | 14024.62 | 5.1 |
| | | SRR4051960<br>SRR4051961 | IMCG47 | 28440.14 | 9.08 |
| | | SRR4051919<br>SRR4101817 | IMCG311 | 15150.43 | 5.29 |
| | | SRR4051941<br>SRR4101820 | IMCG35 | 13081.32 | 5.05 |
| | | SRR4101821<br>SRR4051947 | IMCG36 | 11200.72 | 4.55 |
| | | SRR4101735<br>SRR4052046 | IMCG101 | 13212.90 | 4.64 |
| | | SRR4101737<br>SRR4052247 | IMCG29 | 19661.99 | 6.95 |
| | | SRR4052353<br>SRR4101744 | IMCG04 | 21710.11 | 7.55 |
| | | SRR4101745<br>SRR4052354 | IMCG051 | 16082.35 | 4.36 |
| | | SRR4101749<br>SRR4052120 | IMCG141 | 15530.00 | 5.54 |

| | | | | | |
|---|---|---|---|---|---|
| | | SRR4052177 SRR4101750 | IMCG19 | 18957.64 | 6.5 |
| | | SRR4101751 SRR4052202 | IMCG213 | 13769.74 | 5.3 |
| | | SRR4101747 SRR4052075 | IMCG121 | 12505.18 | 4.68 |
| | | SRR4101753 SRR4052224 | IMCG27 | 15395.76 | 5.75 |
| | | SRR4052234 SRR4101754 | IMCG28 | 16367.42 | 6.06 |
| | | SRR4051772 SRR4096722 | IMCG10 | 36865.38 | 11.57 |
| | | SRR4051866 SRR4101802 | IMCG13 | 28479.47 | 8.98 |
| | | SRR4051917 SRR4101816 | IMCG11 | 24991.59 | 12.38 |
| | | SRR4051928 SRR4101818 | IMCG14 | 31556.88 | 10.34 |
| | | SRR4051939 SRR4101819 | IMCG15 | 24113.19 | 8.04 |
| | | SRR4051964 SRR4101808 | IMCG12 | 39729.48 | 12.8 |
| | | SRR4052103 SRR4101748 | IMCG313 | 30156.49 | 9.86 |
| | | SRR4052272 SRR4101738 | IMCG01 | 26447.13 | 8.33 |
| | | SRR4052346 SRR4101741 | IMCG33 | 29215.81 | 9.71 |
| | | SRR4052349 SRR4101742 | IMCG55 | 36349.01 | 11.9 |
| | | SRR4052525 SRR4096703 | IMCG111 | 37597.81 | 8.58 |
| | | SRR4052590 SRR4096725 | IMCG31 | 28631.05 | 9.39 |
| | | SRR4053363 SRR4096704 | IMCG710 | 23137.77 | 7.63 |
| LNC | Gaizhou, Liaoning | SRR4064255 SRR4107102 | LNC18 | 19120.07 | 7.23 |
| | | SRR4064259 SRR4107114 | LNC21 | 18442.79 | 6.96 |
| | | SRR4067780 SRR4107116 | LNC24 | 25718.73 | 9.33 |
| | | SRR4067798 | LNC33 | 25739.53 | 9.34 |

| | | | | | |
|---|---|---|---|---|---|
| | | SRR4107119 | | | |
| | | SRR4068049<br>SRR4107104 | LNC37 | 22317.17 | 7.98 |
| | | SRR4068051<br>SRR4107105 | LNC38 | 17313.81 | 6.45 |
| | | SRR4068053<br>SRR4107107 | LNC47 | 26501.94 | 9.35 |
| | | SRR4068056<br>SRR4107110 | LNC52 | 25383.53 | 9.16 |
| | | SRR4068057<br>SRR4107111 | LNC54 | 22658.95 | 8.16 |
| | | SRR4068058<br>SRR4107112 | LNC56 | 24913.93 | 8.91 |
| | | SRR4067779<br>SRR4107115 | LNC22 | 11526.85 | 4.75 |
| | | SRR4067781<br>SRR4107117 | LNC31 | 14464.54 | 5.68 |
| | | SRR4067799<br>SRR4107120 | LNC34 | 13730.86 | 5.37 |
| | | SRR4067800<br>SRR4107121 | LNC35 | 12068.03 | 4.93 |
| | | SRR4067801<br>SRR4107122 | LNC36 | 10564.92 | 4.46 |
| | | SRR4068052<br>SRR4107106 | LNC39 | 15629.86 | 5.53 |
| | | SRR4068054<br>SRR4107108 | LNC48 | 11129.55 | 4.47 |
| KO | Korean | SRR1265926 | KOG06 | 45009.41 | 14.92 |
| | | SRR1265927 | KOG07 | 44747.49 | 14.91 |
| | | SRR1265928 | KOG08 | 45333.81 | 14.94 |
| | | SRR1265929 | KOG09 | 44544.50 | 14.76 |
| | | SRR1265930 | KOG10 | 43077.48 | 14.39 |
| | | SRR1265931 | KOG11 | 45720.98 | 15.29 |
| | | SRR1265932 | KOG12 | 44948.16 | 15.11 |
| | | SRR1265933 | KOG13 | 46550.37 | 15.46 |
| | | SRR1265934 | KOG14 | 47704.28 | 15.73 |
| | | SRR1265935 | KOG15 | 45500.75 | 15.26 |
| | | SRR1265936 | KOG16 | 43953.58 | 14.59 |
| | | SRR1265937 | KOG17 | 43054.14 | 14.38 |
| ANG | Evry, France;South Africa;Madagascar | ERR4133570 | FANG0001 | 41597.54 | 13.93 |
| | | ERR4133551 | FANG0002 | 35697.88 | 11.49 |
| | | ERR4133594 | FANG0003 | 18590.68 | 6.53 |
| | | ERR4133600 | FANG0004 | 33849.04 | 11.46 |

| | | | |
|---|---|---|---|
| ERR4133602 | FANG0005 | 31924.81 | 10.83 |
| ERR4133549 | FANG0006 | 37378.87 | 10.28 |
| ERR4133646 | FANG0007 | 34118.43 | 11.55 |
| ERR4133649 | FANG0008 | 29679.00 | 10.08 |
| ERR4133643 | FANG0009 | 37437.20 | 12.86 |
| ERR4133597 | FANG0010 | 16980.35 | 6.01 |
| ERR4133588 | FANG0011 | 39509.40 | 13.07 |
| ERR4133619 | FANG0012 | 18814.75 | 6.54 |
| ERR4133615 | FANG0013 | 39342.96 | 13.72 |
| ERR4133622 | FANG0014 | 33621.92 | 11.7 |
| ERR4133550 | FANG0015 | 42170.35 | 13.13 |
| ERR4133591 | FANG0016 | 32655.34 | 11.11 |
| ERR4133618 | FANG0017 | 40490.15 | 13.93 |
| ERR4133606 | FANG0018 | 35121.24 | 11.88 |
| ERR4133625 | FANG0019 | 37295.04 | 13.03 |
| ERR4133628 | FANG0020 | 34410.02 | 12.12 |
| ERR3281402 | MANG0027 | 103932.50 | 29.9 |
| ERR3281403 | MANG0028 | 68239.26 | 22.11 |
| ERR3281404 | MANG0029 | 80376.99 | 25.78 |
| ERR3281405 | MANG0030 | 79312.28 | 25.62 |
| ERR3281406 | MANG0031 | 62641.33 | 19.91 |
| ERR3281407 | MANG0033 | 76354.83 | 21.87 |
| ERR4133525 | ZANG0374 | 41438.84 | 14.24 |
| ERR4133528 | ZANG0388 | 34554.79 | 11.94 |
| ERR4133517 | ZANG0397 | 31028.87 | 10.73 |

Supplementary Table 3 Sequencing information of Wild goats used in this study

| **Abbreviation** | **Sampling location** | SRR | **ID** | **Size_MB** | **depth** |
|---|---|---|---|---|---|
| IRW | Alamout, Iranian | ERR470100 | IRWG01 | 45083.46 | 15.74 |
| | | ERR470104 | IRWG04 | 43442.83 | 15.49 |
| | | ERR470106 | IRWG06 | 45847.73 | 15.35 |
| | | ERR340426 | IRWG26 | 37967.84 | 13.72 |
| | | ERR340328 | IRWG28 | 19454.66 | 7.24 |
| | | ERR340329 | IRWG29 | 17013.47 | 6.22 |
| | | ERR340330 | IRWG30 | 37395.69 | 13.64 |
| | | ERR340331 | IRWG31 | 34581.11 | 12.73 |
| | | ERR340333 | IRWG33 | 19084.62 | 7 |
| | | ERR340334 | IRWG34 | 37587.13 | 13.58 |
| | | ERR340335 | IRWG35 | 19737.88 | 7.26 |
| | | ERR340336 | IRWG36 | 19252.59 | 7.13 |
| | | ERR340338 | IRWG38 | 34183.70 | 12.48 |
| | | ERR340340 | IRWG40 | 36536.11 | 13.22 |
| | | ERR340341 | IRWG41 | 15342.29 | 5.67 |
| | | ERR340342 | IRWG42 | 19017.87 | 7.03 |
| | | ERR340343 | IRWG43 | 21705.31 | 7.97 |
| | | ERR340344 | IRWG44 | 33823.57 | 12.31 |
| | | ERR340345 | IRWG45 | 35354.22 | 12.85 |
| | | ERR340347 | IRWG47 | 37973.28 | 13.76 |
| | | ERR340348 | IRWG48 | 32400.27 | 11.56 |
| BS | Central Asia | SRR5803211 | BS1 | 40343.35 | 14.58 |
| | | SRR5803192 | BS5 | 34274.71 | 12.13 |
| | | SRR5803193 | BS8 | 48985.98 | 17.08 |
| FAG | Central Asia | SRR5803200 | FAG02 | 27401.58 | 9.71 |
| | | SRR5803204 | FAG05 | 90588.81 | 11.5 |
| | | SRR5803207 | FAG04 | 32469.98 | 31.11 |

BS (Capra sibirica)

FAG (Capra falconeri)

Supplementary Table 4 Sequencing platforms for 236 goats of 13 goat breeds.

| **Breed or Species** | **Abbreviation** | **Sequencing Platform** | **Length** | **Sample Size** |
|---|---|---|---|---|
| Chaidamu Cashmere Goat | CDMC | BGISEQ-500 | 100 | 30 |
| Inner Mongolia Cashmere Goat | IMC | Illumina HiSeq2000 | 100 | 37 |
| Ujumqin Cashmere Goat | UC | BGISEQ-500 | 100 | 12 |
| Liaoning Cashmere Goat | LNC | Illumina HiSeq2000 | 100 | 17 |
| Iranian Wild Goat | IRW | Illumina HiSeq2000 | 100 | 21 |
| Matou Goat | MT | BGISEQ-500 | 100 | 10 |
| Korean native goat | KO | Illumina HiSeq2000 | 100 | 12 |
| Jining Grey Goats | JNG | Illumina Hiseq 2500 | 150 | 10 |
| Jintang Black Goat | JTB | BGISEQ-500 | 100 | 10 |
| Guizhou Black Goat | GZB | BGISEQ-500 | 100 | 8 |
| Yunnan Black Bone Goat | YNBB | Illumina Hiseq 2500 | 150 | 28 |
| Angora goat | ANG | Illumina HiSeq 4000 | 150 | 29 |
| Chengdu Brown Goat | CDB | BGISEQ-500 | 100 | 12 |

Supplementary Table 5 Published ancient samples used in this study.

| Sample | Period | Bioproject |
|---|---|---|
| Potterne1 | Bronze Age | PRJEB26011 |
| Qazvin1 | Bronze Age | PRJEB26011 |
| Bulak2 | Bronze Age | PRJEB26011 |
| Kohneh2 | Bronze Age | PRJEB26011 |
| Acem1 | Bronze Age | PRJEB26011 |
| Acem2 | Bronze Age | PRJEB26011 |
| Azer3 | Bronze Age | PRJEB26011 |
| Darre2 | Chalcolithic | PRJEB26011 |
| Fars4 | Chalcolithic | PRJEB26011 |
| Geor2 | IronAge-Medieval | PRJEB26011 |
| Kazbeg1 | IronAge-Medieval | PRJEB26011 |
| Azer4 | IronAge-Medieval | PRJEB26011 |
| Semnan1 | Neolithic | PRJEB26011 |
| Semnan10 | Neolithic | PRJEB26011 |
| Semnan13 | Neolithic | PRJEB26011 |
| Monjukli8 | Neolithic | PRJEB26011 |
| Blagotin3 | Neolithic | PRJEB26011 |
| Lur12 | Neolithic | PRJEB26011 |
| Blagotin1 | Neolithic | PRJEB26011 |
| Blagotin16 | Neolithic | PRJEB26011 |
| Blagotin2 | Neolithic | PRJEB26011 |

Supplementary Table 6 The genotype frequencies of the homozygous 582bp deletion (-/-) at *LHX2* locus for 15 goat breeds, obtained from whole genome sequencing.

| **Breed** | **LHX2(del/del)** | **Size** | **Genotype Frequency** |
|---|---|---|---|
| CDMC | 21 | 30 | 70.00% |
| YNBB | 8 | 28 | 28.57% |
| CDB | 1 | 12 | 8.33% |
| JNG | 1 | 10 | 10.00% |
| GZB | 1 | 8 | 12.50% |
| IMC | 22 | 36 | 61.11% |
| IRW | 17 | 21 | 80.95% |
| JTB | 0 | 10 | 0.00% |
| KO | 4 | 12 | 33.33% |
| LNC | 9 | 17 | 52.94% |
| MT | 4 | 10 | 40.00% |
| UC | 12 | 12 | 100.00% |
| ANG | 7 | 29 | 24.14% |
| BS&FAG | 1 | 6 | 16.66% |
| AG | 7 | 21 | 33.33% |

Supplementary Table 7 The genotype frequencies of the homozygous 504bp deletion (-/-) at FGF5 locus for 15 goat breeds, obtained from whole genome sequencing.

| **Breed** | **FGF5(del/del)** | **Size** | **Genotype Frequency** |
|---|---|---|---|
| CDMC | 23 | 30 | 76.67% |
| YNBB | 0 | 28 | 0% |
| CDB | 0 | 12 | 0% |
| JNG | 2 | 10 | 20% |
| GZB | 0 | 8 | 0% |
| IMC | 30 | 37 | 81.08% |
| IRW | 0 | 21 | 0% |
| JTB | 0 | 10 | 0% |
| KO | 0 | 12 | 0% |
| LNC | 17 | 17 | 100% |
| MT | 0 | 10 | 0% |
| UC | 0 | 12 | 0% |
| ANG | 1 | 29 | 3.45% |
| BS&FAG | 0 | 6 | 0% |
| AG | 0 | 21 | 0% |

Supplementary Table 8 The frequencies of the 582 bp deletion near *LHX2* with PCR amplification.

| Breed | **del/del** | **del/+** | +/+ | **Sum** |
|---|---|---|---|---|
| CDMC | 131(54.13%) | 91(37.6%) | 20(8.27%) | 242 |
| IMC | 8(66.67%) | 3(25%) | 1(8.33%) | 12 |
| UC | 30(75%) | 9(22.5) | 1(2.5%) | 40 |
| JNG | 2(16.67%) | 8(66.67%) | 2(16.66) | 12 |
| WSW | 0 | 2(16.67%) | 10(83.33%) | 12 |
| JTB | 1(8.33%) | 5(41.67) | 6(50%) | 12 |

The frequencies of homozygous (del/del), heterozygous (del/+) and wide type of the 582 bp deletion near *LHX2*, obtained from PCR amplification.

Supplementary Table 9 The frequencies of the 504 bp deletion near *FGF5* with PCR amplification.

| Breed | del/del | del/+ | +/+ | Sum |
|---|---|---|---|---|
| CDMC | 194(80.16%) | 45(18.6) | 3(1.23%) | 242 |
| IMC | 12(100%) | 0 | 0 | 12 |
| UC | 3(7.5%) | 19(47.5%) | 18(45%) | 40 |
| JNG | 3(25%) | 8(66.67%) | 1(8.33%) | 12 |
| WSW | 0 | 1(8.33%) | 11(91.67%) | 12 |
| JTB | 0 | 0 | 12(100%) | 12 |

The frequencies of homozygous (del/del), heterozygous (del/+) and wide type of the 504 bp deletion near *FGF5*, obtained from PCR amplification.

Supplementary Table 10 Primers used for amplification of deletion seences

| Primer names | Sequence 5’-3' | Length (bp) |
|---|---|---|
| LHX2_del_F2 | CAGTACGGAGCAAGTAAACGG | 630 bp for del/del |
| LHX2_del_R2 | ACCATTCCACTTGTCCACCT | 1214 bp for +/+<br>630/1214 bp for del/+ |
| FGF5_del_F1 | ACAGCGTGTGATCTTTTCTCTG | 235 bp for del/del |
| FGF5_del_R2 | TCTTGGTCTGGCTGTGATCA | 742 bp for +/+<br>235/742 bp for del/+ |

Supplementary Table 11 Samples for PSMC analysis

| Sample | depth(X) |
|---|---|
| BBG31107 | 26.94 |
| FANG0017 | 13.93 |
| WMG0004 | 15.93 |
| CMS0088 | 13.21 |
| HXR0127 | 12.6 |
| IRWG01 | 15.74 |
| KOG14 | 15.73 |

#### Supplementary sequence 1

Distal enhancer-like signatures are shown in red, the CTCF binding site is displayed in bold, and the 582bp deletion sequence is underlined.

>goat:chromosome chromosome:ARS1:11:94266769:94278930:1
ATACGAGTTGCCATGACTATTCTGTTGCTTGGATTCATTCACCATCTACTCTCACCAGT
TGTACTAATTAAATCTTCAGAGCTCTTCCAGGATCTCTATCGCTTTCTTTGCTTCAGAA
AGTCAATCAGAGTTTATTGCCCCGAAAAGGTAGGATTCAGAGTCAACATTTACTAAA
TGAGTAGTATGTGTCCAGCAATGGGCTACTGGCTTTCATATCTTTCAAAAACCCTGAG
GGGAGTATTAACACTACTCTCTTCTTACTGAAAAGATTAAGTTGCCTAGGGCAACATG
GCTAGGGAGGATCAAGATACACAGGAAGAATTTCAACAAAACCATTATGACTTGAAG
CCTGGTGCTCTCTCCTCTTGTCCTTTTTCTTGGGAAAAGCTAGCAGAAGCACGGCTGT
CTGAATCATCTAGCACGGAAAACACATAAATATCAAGACAGCACAAGAGAAAAAAC
ATGAATTTTCACAAGGTCTATATGAACTCCTAATGATATAGTCTAACCTCACATCATA
CTTCATTCGCTCATCAGTGACCATGCTTTCAGTTTGCAGAAATAAAAAAAAAATTTGA
GTTCTTGACTTCTGACCACATAGTAGGTGCTCCAATTGTGTTAACAAAATCCACACAG
ATGTCTGCCAATGTGAGTGCCATTAGGGGTTTTTCAAGTATATTAAGATGTCAGTGAC
TTGGAAGGAAGTTATAACCAACCTAGATAGCATATTCAAAAGCAGAGACATTACTTT
GCCAACTAACACCCATCTAGTCAAGGGTGTGGTTTTTCCAGTAGTCATGTATGGATGT
GAGAGTTGGACTGTGAAGAAAGCTGAGTGCCGAAGAATTGATGCTTTTGAACTGTGG
TGTTGGAGAAGACTCTTGAGAGTCCCTTGGACTGCAAGGAGATCCAACCAGTCCATT
CTGAAGGAGATCAGTCCTGGGTGTTCTTTGGAAGGAATGATGCTAAAGCTGAAACTC
CAGTACTTTGGCCACCTCATGCAAAAAGTTGACTCACTGGAAAAGACTCTGATGCTG
GGAGGGGTTGGGAGCAGGAGGAAAAGGGGACGACAGAGGATGAGATGGCTGGATG
GCATCACCGACTCAATGGATGAGAGTTTGAGTGAACTCCGGACAGGGAGGCCTGGCA
TGTTGCAATTCATGGGGTCGCAAAGAGTCGGACAAGACTGAGTGACTGAACTGAACT
GAACTGACTTGAATATCACCTCTTAGTGTTTTTTGGTAGAGATCTGATTCTTAAAAGA
GTCATATAAAAAAGTTATCAATCCTATAATCTAAGAAACTGACTACACCACTAAAAT
AATACACAAGATACTCTGCCAATGTTCTCAATGCAATGGCTGAAAAGAAACATCACC
TCTCTAGATCCATTTATACAGATTTATTTAAAACTACCCTAGTGGACTACATCAGCAT
CCAATGGTACTGAGAGCTTAAATTACTCTTAGCTCTGTCCTGTTCTAGACTGTTGTTGC
TGCTGCTGCTAAGTTGCTTCAGTCATGTCTGATTCTTTGCGACCCCATAGACTCAGAG
CTCTTTCAGAAATATGGAAATAATTTTTAGTGCTAATCTGTACTCACAAATGGAGAAG
GAAATGGCAACCCACTCCAGAATTCTTGCTTAGAGAATACTGTGGACGGAGGAGCCT
GGTGGGCTGCTGTCCATAGGGCTGCACAGAGTCGGACACGACTGAAGCGACTTAGCA
CGCATGCATGCACCGGAGAAGGAAATGGCAGCCCACTCCAGTGTTCTTGACTGGATA
ATCCCAGGGATGGTGGAGCCTGGTGCGCTGCTGTCTATGGGGTCGCACAAAGTCGGA
CACGACTGAAGCAACTTGGCAGCAGCAGCAGCAGTACTCACAAATGCCTTCTAAACA
AACTTAGTCAGCCTTTATTATATAGTCTCAAGATGAAAAGGACTCTATTGATGCAAAA
TAATCAAAGAATAAATACTGTCTTAGAGAACTGGTCCTGAATCTGGTTATGTATTAGT
AATTTGGGAAACCTTAAAAAAATAGACATCCCACAGTCCCATCTTAGACTTTTTGAAT
TATTATTATAATAAAAATTTAAGTTCTTGGGTACTACCTCTACACATTCTGATTCAAAA

GGTCTAATTTTTATTTAAATTGGAGAATAATTAGTTTACAACATTGTGATGTTTTTAGC
CATACATCAACATGAATCAGCCACAGGTATACATTTGTCCCACCCCATCCTGAATCCC
CTCCCACCTCCTTCCTTACCCTATTCCTCTGCGGTTGTCCCAGAGCATCAGCTTTGAGT
GCCTTGCTTCATGCATCAAACTTGCACTGGTCATCTATTTTACACATAGTAATGTACAT
GTTTCAGTGCTATTGTCTCAAATCATCCCCTGCTCTCCTCCTCTCCCTCTGAGTCCAAA
AGTTTGTTCTTTACATCTGTGTCTCCTTTACTGCCCTGCATGTAGGATCATTGGTATTG
TCTTTCTAAATTCCATATATATGCGTTAATATATAGTATTTGTCTTTCTTTTCAACTTAT
TTCACTCTGTATAATAGGCTCCAGGTTCATCTACCTCATTAGAACTGACTCAAATGCA
TTCCTTTTTGTAGCTGAGAACTTAAATTTTTAATAAGCTCTCTGAGTGATTTCTAACAC
AACTAACTCAGGGATCAGCATTTGGGAATCACTAACAATCTGAATCCTCTGTCTAGTC
CAAAACATGTGCTCAATGTGTTGAAGCAAGAAGGGCAGGAGAGGAAGGTAAGAACT
AGTAGGGAGACCGTGATTGGTTGAAATCACAGCAATATTTAAAGTACTACAAATTAA
AGTGTTGTTACTCTATATATTATATAGCTTATACATTCTCTATAACATCAATGTTCTTC
ATTCTTATAGTCTAACATGAACAAGAGAAAAGGAACTCGCACATAATTAAATGAAGA
ACATAGTTCAAAAAGCAGCCAAGTGTATTTACATTTAAAGACCAGAGAAAAACCACA
TAAATAAAACATACCAATCTTTTCTATTTCCTGCCAATGGAAGTTCCTTTCATTTATTT
CTAACAGTTTTTAATATTCTCTGAGCCACAATAAAATTTTATGTGATGGAGATTAGAG
AGGCTATGCTATTTGAAAGTTTGGCTGCAGAGTTTGTATTTCATGCCACTTCATTACAT
TTTGTTAAGGATTTCCCTCTTCACCAAAAGGGATTAGTTTTCTTCTCACTTCTTTGAAT
ACTCTTCTTAGGTTGTTAGTGTGAACTAGGTTAATCTCTCTTTTAACATATAAAAGATG
AACAAACAAAATGTTCAATTTATTTTATTTTTTTAAATGGAACACTAAATTTAGTCTTA
GAATTCTCTGTTTGGTAAGTTTCCAGTTACAAGGTCTTGGCTTCCCTTGAGCACAGTTT
GAAACAAAATAATGCAGTTTGTTCTGATGGCTGGAACATGCAGAGTGGAGTGTGCAG
GCAAAGAGGCTCTGGCTCTGGTCCAGTATTTCAAACCTTCTTAAACTGCCACTCTCAT
CTAGGCTTCAAGGGCTCCAGATAAGCAAACAGGATTAGTTATTTCTAGCCCAGAAGA
GCTAGAAAAGCTTTATACACATCATTTAGGGACATAGCCCACTCTACAAGTCAGGGT
AGTCATGACCAGGTAGATCTGATCAGGAAAGAAGTAAAGATGAAGCTAAAGATCTG
AAACATTCCCTAGTCCACAAACTGACTCTTCTTAGGTCACAAAACCAGGTGGTGCTAA
GACACTGGATGCTGTGAATGGAAGTGGATAGGACAGGACTGGAGCAAGGTTGGGAA
GTTCCCCATCCCTTCATGCCAACACTCAAGTGTCTACTAAGGCCACAACTGAGGGTGC
TGGGGGGAAAAAGAAATACCACCCACCACTCTCCAGGCCATCAAGTCCTGCCCAAGG
CACATTTCTGGGACTAATTGGAAGAAATTGACGGAGAGATATTGGGCTTTTTTTTTTT
TTTTTTCCTTCTCTACATGGTGTATTTGGAGAAGGAAACAGCAACCCACTCCAGTATG
CTTGCCTGGAAAATTTCATGGACAGGGAGCCTGGTGGGCTCTAGTCCATGGGGTCAC
AGAGTCAGACACGACTGAGTGATTAAATGACATAACATACTATTAGAAAAATAGTAA
TTCTTTTCATAGTTGTGGGTATCACACAAATAATTAGGTGTGGAGGCTCAATTAATTA
GGTAACAGCTTTGCCACCAAATTGTGCTTCCTTCCTAAGAACTTGTTTTAAAAAGCAT
CTCTTAAGTGTGACAGATATTCTTAACAGTGTGCCTCTGGTTCTCGTGACAGTTCCCA
CACAAGGATTCTGATGTTTGTTTGAGAGGTGACACCCAATCAGTGTAACCTGGTCCCC
ACAGCAAGCTCTTTGAAGGGGCCAAGATTCATCTGTGTAACTCCTGGTTGTTACTCCC
TTTGCTGAAGCCAAAGCCTCCCTTCCTCTGACACATCATCATAATGACCCACCTGCAT
GTCCTTTTCTTCTATTACGAGGGCATAACCTTGGGCAAGAGAACCCATCTCTCTGAGT
GGATGGATCATTCTGTAAAATGGGGATAATATTGCTCTTCTAGGAATTCTGTTGCAAT
TAGATGATATTATTTCTATAAAGGTTTTATTTTAGACCTACTTGCTATCCACAAGAGTA

ACCAAAAAACAATCTGAGTTAAACTCAGATTTGAAAATCTGGAAGAAAAATGGGGCC
TTTGTGCTAGAAGAAATCTCACTGAAGCATTTACATTAGAGAACAAGATGTAGTTAA
ATCTTCCCTATTTATCATTTATAAAGGAAAACAATCAATATACACTGCACAAAAAGAA
CAAAAAGTTAACATTTTTCTTACACAGAATATCACAACATTATTGACCGGGAAAGCCT
**GAAATGAATATTACCTGACTCTGAACTAAACATGGCTGGGTTTGTCTCTATTGAC**
**ACACAGTCAGTTTTATGAAACATGGCACTCTGGGATGTTCAACAGCTTTCATCTT**
**TCAGATGCTTGATGGCTCTGAGAAGCATCAAGCCTACATCTTTATGAAACAAAAT**
**TGAGAAAAATGTGTTTTGTCAAGGTATGGAAGGCCACATCCGGATCTGTCAAGA**
**AAGGGTCATCAAAAACTTCAGGCTACAGTATTCTTTGCATTTTCCTTCATCAAAA**
**AGAAAATAAAAAGTCAGTACGGAGCAAGTAAACGGTAACAATTAGAAAGCTTTT**
**A**AACTCTAAAATTGAAGAGGTCCAAAGGCTTGGATAAAAGGGATGTCTACTCAGTTA
CTTATCCCCAGAGAAACAACACAAAACAGATCAGGGTGTTGCTGATCCCCTATGAGG
TAAAAGAAGACCTTCTCCTTCCCTAAAGAGATATCCACCATCAATTAGTAAAACTTAG
AACTCAGTTAAGAATCAACTGTCTGTTTAAAAGTTTTTAAGTCAAAGTCATGTATTTA
TAGCCTGAAAAACAAAACAAAAAGTGTTAAAGGCTTATACTACTGACAAATAGCAGC
CCATCTCATGATATCCCTTACCTTCAATCAAAAGTAATCACATTCAACACAATATTTA
TCTCCTTATTTTTAAAAAGTATTTAATTTAAAAATTTAAACATTAATCTTTGACTTCCC
ATTATAGATATTAATAATTTATCAAGAACATATACATATACCTTATGCCAGACTGTTC
TAGGAGTTTTGCAACTCGTTTTATTTGACATGAGGTCACTGGTGACCAAACAAAAACT
GTTTCAGTGGAGGCAGGGGCTAAATCACAACACAAGACAGAGGAGAGAATGAGAAG
AGAGGAATTAGACAGTAAACAGAAGCAACTCTCTTGAGGAATTGCATTCTAAAAGGA
AGAAAAGAAATGGGATAAAAGAGATGTGTTTAAGGATGTTTTGTTTTGAAGTTTGGG
AGACACTATACTGTGCTAGTAAGTGCATAAGACTGTTCAGTTCAGTCGCTCAGTTGTG
TCCGACTCTTTGCGACCCCATGAATTACAGCACGCCAGGCCTGTACCAGTAAAGAAA
ACAAAAAACAAAAAATGATGCAGGAGAGAGAAAGCTCAAGGCTGGAGGAATTCCGT
AAGGTGACAAGAAGGATGCTGCAGTGTTAGCTTTCCCACAAGGCACTCCTTGGTCAT
CTAGCTAAACTGCAATCCCTGCTACACCAGTCGCCATCCACTCTCCTCCTCTGCTTAAT
TGTACTCCTTGTACTTAGAACTAGCTACTATTCTATATTTTCAACCCACCTCATTTATT
GATCTACTCGACTAGAACATAAGCTTCATATGAGGACAGGGACTTTTGCCTATTTTGT
TCACTGCTGTATCACCAGTACCTATGACAGTGCCACAAAAACTATTTGCTGAATGAAG
TCAGGTGGACAAGTGGAATGGTTGGCCTAGACAGAATGGACAGTTCATCCACAATGG
AGAAGGTGAGTGGAGGACTAGCTACCGCAGGTTGACAGGTGTGGTAGGGGCTGGTGC
CACTCTCTCCTGAGCGCTTTCATTAGATAAAGATGGAATTTTAGGTAAGAAATATTTT
TCCCTTAGAAGTTTAGGCACTGCTTCACTATCTTCTCGTTTCTAGAACTGCCACTGCGA
AATGTTGTGCAAGTCTGATTCTTAATGTTTATACAATGACTACTTTGTCTCTATCTCTG
TATTGTAAAATTACATGATAAAAAGGGCTTTTTAACTTTCAAAGATTTCCTGAAGCCA
AAACCTCCCTTCCCCTGACACTACATCATAACGACCAATGTGCCTGTCTCTTCTATGA
CTACCACGTGGCCTTAGGCAAGTTAACCCATGTATCTGAGTGAGTGGGCCATTCTGTA
AAATGGGGATAATATTGCTTTTTAGACTTTTAAAGAATTCTGAA

#### Supplementary sequence 2

Ins sequence(551bp)

GAAAAACAAAACAAAAAGTGTTAAAGGCTTATACTACTGACAAATAGCAGCCCATCTC
ATGATATCCCTTACCTTCAATCAAAAGTAATCACATTCAACACAATATTTATCTCCTTATT
TTTAAAAAGTATTTAATTTAAAAATTTAAACATTAATCTTTGACTTCCCATTATAGATATT
AATAATTTATCAAGAACATATACATATACCTTATGCCAGACTGTTCTAGGAGTTTTGCAA
CTCGTTTTATTTGACATGAGGTCACTGGTGACCAAACAAAAACTGTTTCAGTGGAGGC
AGGGGCTAAATCACAACACAAGACAGAGGAGAGAATGAGAAGAGAGGAATTAGACA
GTAAACAGAAGCAACTCTCTTGAGGAATTGCATTCTAAAAGGAAGAAAAGAAATGGG
ATAAAAGAGATGTGTTTAAGGATGTTTTGTTTTGAAGTTTGGGAGACACTATACTGTGC
TAGTAAGTGCATAAGACTGTTCAGTTCAGTCGCTCAGTTGTGTCCGACTCTTTGCGACC
CCATGAATTACAGCACGCCAGG
